# Conjunctival epithelial cells resist productive SARS-CoV-2 infection

**DOI:** 10.1101/2021.12.20.473523

**Authors:** Robert M Jackson, Catherine F Hatton, Jarmila Stremenova Spegarova, Maria Georgiou, Joseph Collin, Emily Stephenson, Bernard Verdon, Iram J Haq, Rafiqul Hussain, Jonathan M Coxhead, Hardeep-Singh Mudhar, Bart Wagner, Megan Hasoon, Tracey Davey, Paul Rooney, C.M. Anjam Khan, Chris Ward, Malcolm Brodlie, Muzlifah Haniffa, Sophie Hambleton, Lyle Armstrong, Francisco Figueiredo, Rachel Queen, Christopher J A Duncan, Majlinda Lako

**Author notes:** joint first authors (Robert M Jackson, Catherine F Hatton, Jarmila Stremenova Spegarova, Maria Georgiou). joint corresponding authors, *Correspondence should be addressed to:* Dr. Rachel Queen, Dr. Christopher Duncan and Prof. Majlinda Lako.

## Abstract

Although tropism of SARS-CoV-2 for respiratory tract epithelial cells is well established, an open question is whether the conjunctival epithelium is also a target for SARS-CoV-2. Conjunctival epithelial cells, which express viral entry receptors ACE2 and TMPRSS2, constitute the largest exposed epithelium of the ocular surface tissue, and may represent a relevant viral entry route. To address this question, we generated an organotypic air-liquid-interface model of conjunctival epithelium, composed of progenitor, basal and superficial epithelial cells and fibroblasts, which could be maintained successfully up to day 75 of differentiation. Using single-cell RNA Seq, with complementary imaging and virological assays, we observed that while all conjunctival cell types were permissive to SARS-CoV-2 genome expression, a productive infection did not ensue. The early innate immune response to SARS-CoV-2 infection in conjunctival cells was characterised by a robust autocrine and paracrine NF-Kβ activity, without activation of antiviral interferon signalling. Collectively, these data enrich our understanding of SARS-CoV-2 infection at the human ocular surface, with potential implications for the design of preventive strategies and conjunctival transplants.

## Introduction

Coronavirus disease (COVID-19) is an infectious disease caused by the severe acute respiratory syndrome coronavirus (SARS-CoV-2). Since its emergence in late 2019, 270,510,989 infections and 5,324,329 deaths have been reported up to 13^th^ December 2021 (WHO, 2021). It is well established that cells of the nasal and respiratory epithelium are the principal targets for SARS-CoV-2 (Sridhar and Nicholls, 2021). This tropism is a central factor in transmission and pathogenesis of COVID-19. Viral tropism for other cell types has also been reported, including neurones (Ramani et al., 2020, Song et al., 2021) and gut epithelial cells (Lamers et al., 2020), likely contributing to other aspects of COVID-19 pathogenesis. The ocular surface is a defined route of entry of several viral pathogens (Belser et al., 2013, Chu and Pavan-Langston, 1979). An important unresolved question is whether SARS-CoV-2 is similarly capable of infecting cells of the eye surface. Beyond the obvious importance of this question to our understanding of viral pathogenesis, it has implications for the use of personal protective equipment (PPE), such as visors or other forms of eye protection, in healthcare settings.

Host cell receptor expression is a major determinant of viral tropism. The SARS-CoV-2 spike protein binds angiotensin-converting enzyme 2 (ACE2) enabling viral entry; spike-mediated membrane fusion is facilitated by the host transmembrane protease serine type 2 (TMPRSS2) (Hoffmann et al., 2020). Work from our group and others has demonstrated expression of key entry receptors by cells of the ocular surface (Collin et al., 2021b), suggesting that the eye may be a plausible route of viral entry. The ocular surface comprises superficial conjunctiva, limbal and corneal epithelium and associated glands. Together with the tear film, these cell types form the first line of defence against infection of the eye (Fleiszig et al., 2002). Evidence for ocular tropism of SARS-CoV-2 remains inconclusive, as recently reviewed (Armstrong et al., 2021). Clinical reports suggest SARS-CoV-2 can be detected in tears and/or conjunctival swabs from COVID-19 patients, although the percentage of patients with detectable viral RNA was low (0-5.3%) (Zhong et al., 2021, Seah et al., 2020). Clinical syndromes of ocular infection (e.g., conjunctivitis, keratoconjunctivitis, etc) are also infrequently reported (Armstrong et al., 2021) in patient cohorts, although they are noted in case reports (Ozturker, 2021, Scalinci and Trovato Battagliola, 2020). Consistent with the relatively low frequency of detection of SARS-CoV-2 in clinical ocular specimens, a post-mortem study identified SARS-CoV-2 RNA in ~ 13% of 132 post-mortem ocular tissues from 33 infected patients (Sawant et al., 2021). Conversely, in another post-mortem study, viral protein was detected by immunofluorescence analysis in 3/3 patient ocular tissues analysed, with positive staining found mainly in the limbus, and the central cornea exhibiting very low levels of viral detection (Eriksen et al., 2021). The ocular route appears to be a bona fide route of SARS-CoV-2 transmission in studies of rhesus macaques and Syrian golden hamsters (Imai et al., 2020, Deng et al., 2020, Hoagland et al., 2021). Ocular inoculation with SARS-CoV-2 resulted in a mild lung infection in these models, however evidence for direct infection of the ocular surface was not sought (Deng et al., 2020). Notably, the nasolacrimal duct connects the ocular surface to the nasal mucosa, providing indirect access to nasal mucosal tissues from virus inoculated at the ocular surface; thus, ocular transmission does not necessarily imply permissiveness of the ocular surface.

Other studies have directly assessed the capacity of human ocular cells or tissues to support experimental infection *in vitro*. Miner and colleagues reported that human corneal cultures were resistant to SARS-CoV-2 infection (Miner et al., 2020). This resistance was not mediated by an innate antiviral type III interferon (IFN) response, as was the case for other viruses studied, suggesting alternate mechanism(s) of SARS-CoV-2 restriction. In compatible findings using cultured corneal, limbal, scleral, iris, retinal and choroid cells from healthy cadaveric human donor eyes, alongside an induced pluripotent stem cell organoid system, Erikson and colleagues identified limbal cells to be more permissive than corneal cells to SARS-CoV-2 infection (Eriksen et al., 2021). Consistent with this, Sasamoto and colleagues showed that limbal cells express high levels of ACE2 and TMPRSS2 (Sasamoto et al., 2021). The main limitation of the studies described above was that the conjunctiva, which occupies the largest ocular surface area and contains cells expressing *ACE2* and *TMPRSS2* (Collin et al., 2021b), was not investigated. To our knowledge, one recent study has assessed the permissiveness of conjunctival cells. In this study, Singh et al dissected conjunctival cells and infected them under submerged culture conditions (Singh et al., 2021). They reported detection of viral RNA expression alongside expression of innate inflammatory mediators, suggestive of infection. They also detected spike protein in the superficial conjunctiva of patients that had succumbed to COVID-19. In cultures, the expression of viral protein declined rapidly from 24 to 72 hours post-infection, suggesting that conjunctival cells may be unable to sustain infection. However the capacity of these cells to support a productive infection, via assembly of nascent viral particles or release of infectious virus, and the responses of individual conjunctival cell subtypes (e.g. superficial or basal conjunctival epithelial cells and epithelial progenitor cells) were not formally assessed. Further studies are required to determine whether the conjunctiva is a permissive tissue and might act as an entry portal for SARS-CoV-2.

Currently there are limited *in vitro* cellular models available for modelling the human conjunctiva. Garíca-Posadas *et al*. developed two three-dimensional fibrin scaffolds on which they could seed human conjunctival epithelial cells (Garcia-Posadas et al., 2017). These models maintained their epithelial-like properties for 14 days before epithelial mesenchymal transition began, resulting in loss of MUC5AC expression by day fourteen. A slightly different approach was taken by Chung et al. to generate a multi-layered construct replicating the conjunctiva (Chung et al., 2007), comprised of a 6-8 layered epithelium with a high proportion of goblet cells. These constructs were characterised by the secretions of the membrane bound MUC1, MUC4 and MUC16 and the secreted MUC5AC between 1-3 weeks of culture. The relative limitation was that replicative senescence was reached after 3 weeks and cells started to detach from the construct. An alternative model was developed to generate progenitor cells for use in transplantation. Conjunctival epithelial cells derived from patients were grown at the air-liquid interface (ALI) and after two weeks of differentiation expression of MUC5AC, KRT3, KRT19 and KRT12 could be detected by immunofluorescence analysis (Jeon et al., 2013). This tissue was used for transplantation in animal models and consequently longevity of the culture remains to be assessed.

To overcome these issues and address definitively the permissiveness of conjunctival epithelium to SARS-CoV-2, we describe the generation and full characterisation of an ALI organotypic conjunctival epithelial model, composed of progenitor, basal and superficial epithelial cells and fibroblasts. Using single cell RNA-Seq, with complementary imaging and virological assays, we define the cellular permissiveness of various epithelial cell types in the conjuctiva and determine the cell type specific innate immune response to SARS-CoV-2 infection.

## Results

### Generation and characterisation of the ALI conjunctival epithelium model

SARS-CoV-2 entry factors *ACE2* and *TMPRSS2* are co-expressed at the highest level in the superficial conjunctival epithelium, which led us to hypothesise that ocular surface epithelium may provide an entry portal for the virus (Collin et al., 2021b, Sungnak et al., 2020). To test this hypothesis, we developed a protocol to generate an ALI conjunctival epithelium model, containing several epithelial cell types akin to human conjunctiva *in vivo*. To this end, epithelial progenitor cells from the perilimbal conjunctival epithelium were *ex vivo* expanded on mitotically inactivated 3T3 feeder cells (Spurr-Michaud and Gipson, 2013) (**Figure 1A**) and then matured at an air-liquid interface (ALI) to induce differentiation. Conjunctival epithelium shares similar characteristics to airway epithelial cells (Chung et al., 2007), hence similar culture conditions that promote the development of an upper airway ALI model were applied during ALI differentiation as outlined in the methods section and **Figure 1A** (Djidrovski et al., 2021).

**Figure 1:**
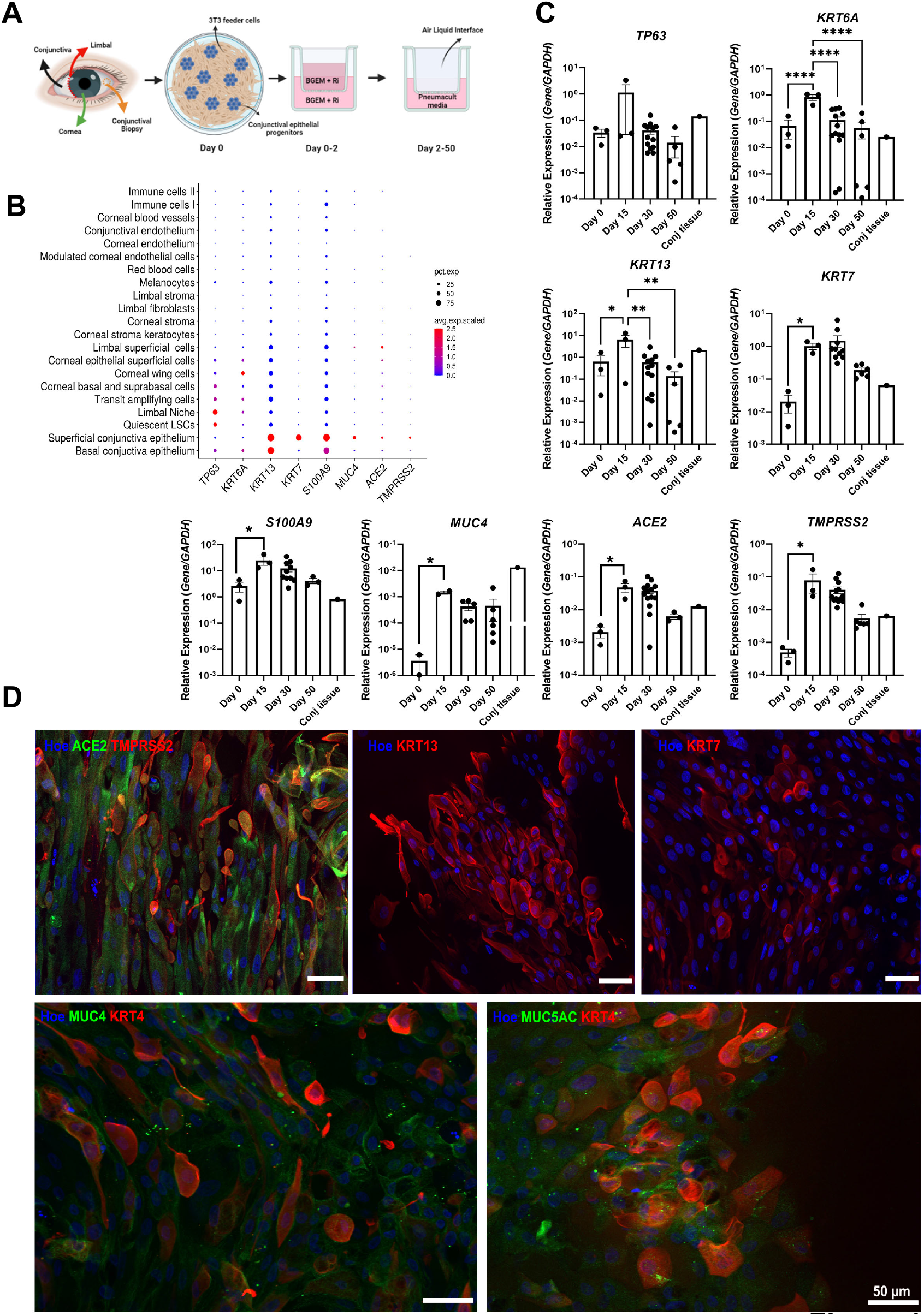
Generation and characterisation of the ALI conjunctival organotypic culture model. **A**) Schematic summary showing the key steps involved in generation of the ALI conjunctival organotypic culture model. **B**) RNA expression of progenitor, basal and superficial epithelial cell markers in specific cell types in the human adult cornea and conjunctiva. Single cell RNA-Seq data from Collin et al. (2021a) were used for this analysis. Raw expression values were normalised, log transformed and summarised. The size of the dots indicates the proportion of cells, while the colour indicates the mean expression. **C**) Quantitative RT-PCR analysis showing expression of progenitor and basal epithelial marker (*TP63*), basal and superficial epithelial cell markers (*KRT6A, KRT13, KRT7, S100A9, MUC4*) and SARS-CoV-2 entry factors (*ACE2* and *TMPRSS2*) in ALI conjunctival organotypic culture model and primary conjunctival tissue. Data shown as mean ± SEM, n=3-14 experimental repeats in three different donors, * p < 0.05, ** p < 0.01, *** p < 0.001, one way ANOVA with Tukey’s multiple comparisons. **D**) Immunofluorescence analysis showing expression of conjunctival epithelial marker (KRT13), superficial conjunctival epithelial marker (KRT4, KRT7), goblet cell marker (MUC5AC), mucin producing cells (MUC4) and SARS-CoV-2 entry factors (ACE2, TMPRSS2) in day 15 ALI conjunctival organotypic culture models (representative of repeat experiments in three different donor conjunctival ALI cultures). Hoe-Hoescht.

Samples were acquired at day 0, 15, 30 and 50 of differentiation and analysed by quantitative RT-PCR for expression of several markers characterising epithelial progenitors and conjunctival basal epithelium (*TP63*), conjunctival epithelium (*KRT13, S100A9*), superficial (*KRT7, MUC4*) and basal conjunctival epithelium (*KRT6A*) (**Figure 1B**). This analysis showed the persistence of epithelial progenitor cell marker *TP63* throughout the differentiation period and a significant increase in the expression of *KRT13* and *S100A9* from day 0 to day 15 (**Figure 1C**), indicating the onset of differentiation towards conjunctival epithelium. Notably, the expression of *KRT6A* and *KRT7* was increased from day 0 to day 15 of differentiation, indicating specification to both basal and superficial conjunctival lineages respectively. The expression of superficial conjunctival epithelial markers (*KRT7, MUC4*) was maintained during the differentiation process, while those of basal conjunctival epithelium (*KRT6A*) declined, suggesting that the culture conditions were more permissive for the development of superficial conjunctival epithelium. The expression of *ACE2* and *TMPRSS2* increased significantly during the first 15 days of differentiation at ALI and was maintained at similar levels to the uncultured conjunctival tissue (**Figure 1C**). These findings are consistent with reports of increased *ACE2* expression by respiratory epithelial cells during maturation at ALI (Sungnak et al., 2020), and were corroborated by immunofluorescence (IF) analysis, showing co-expression of ACE2 and TMPRSS2, as well as abundant expression of KRT13, and superficial conjunctival epithelium markers KRT4 and KRT7 at both day 15 and day 30 of differentiation (**Figure 1D, Figure 2A**). Importantly, we were able to detect expression of MUC5AC, indicating the presence of goblet cells in our ALI conjunctival epithelium culture model (**Figure 1D, Figure 2A).**

**Figure 2.**
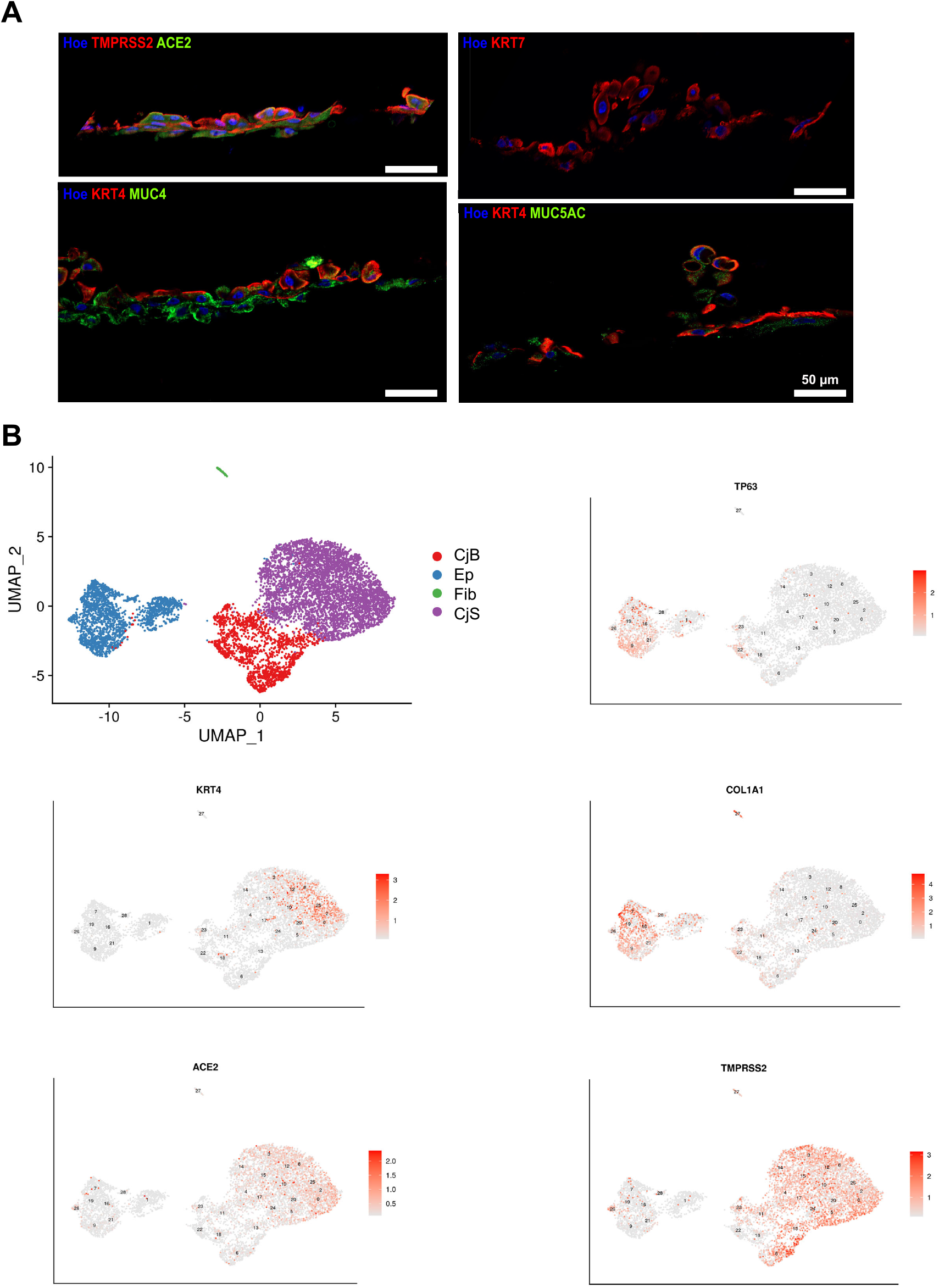
Characterisation of ALI conjunctival organotypic culture model at day 30 of differentiation by immunofluorescence and single cell RNA-Seq. **A)** Immunofluorescence analysis showing co-expression of ACE2 and TMPRSS2 in the superficial layer of the ALI conjunctival organotypic model. KRT7 and KRT4 were predominantly located in the superficial layer, while MUC4 was detected throughout (representative of repeat experiments in three different donor conjunctival ALI cultures). Hoe-Hoescht. **B**) UMAP visualisation of scRNA-Seq data from conjunctival ALI cultures (8,202 cells from three different donors) showing the presence of epithelial progenitors (EP), superficial (CjS) and basal conjunctival epithelium (CjB) and fibroblasts (Fib). Expression of progenitor and basal epithelial marker *TP63*, superficial conjunctival marker *KRT4*, fibroblast and progenitor marker *COL1A1*, and SARS-CoV-2 entry factors, *ACE2* and *TMPRSS2* are shown as superimposed single gene expression plots on the UMAP.

Single cell RNA-Seq analysis of day 30 ALI samples obtained from three different donors was performed revealing the presence of a predominant superficial conjunctival epithelial cluster comprising 55.4% of the total cells analysed (**Figure 2B**) as well three other smaller clusters defined as basal conjunctival epithelium, epithelial progenitors and fibroblasts (comprising 19.2%, 24.7% and 0.8% of the total cells analysed respectively) based on highly expressed markers (**Table S1**). Notably, the expression of *ACE2* was highest in the superficial conjunctival epithelium (**Figure 2B**), whilst *TMPRSS2* was expressed at high level in the superficial but also some basal conjunctival epithelial cells. In total, 26% of cells co-expressed *ACE2* and *TMPRSS2*. To assess the representativeness of the ALI conjunctival model, we correlated expression of the top 2000 highly variable genes between each organoid cell type and the twenty-one-adult cornea-conjunctival single cell RNA-Seq clusters reported earlier this year by our group (Collin et al., 2021a). The strongest correlations were observed between the ALI superficial conjunctival epithelial cells to superficial conjunctival epithelium *in vivo* (correlation coefficient 0.67) and ALI basal conjunctival epithelial cells to basal conjunctival epithelium *in vivo* (correlation coefficient 0.58).

Immunofluorescence analysis at day 75 of differentiation revealed the presence of a multi-layered epithelium, characterised by predominant apical expression of the superficial conjunctival markers KRT7 and KRT4 and widespread expression of MUC4 (**Figure S1A)**. Single cell RNA-Seq (**Table S1**) at this later time point revealed the presence of similar cell clusters to day 30 as well as superficial cell expression of *ACE2* and broader expression of *TMPRSS2* (**Figure S1B**). By immunofluorescence analysis, cells co-expressing both ACE2 and TMPRSS2 were found both on the apical layer and inside the ALI cultures (**Figure S1A**): those comprised 36% of the total cells analysed by single cell RNA-seq.

Transmission electron microscopy (TEM) analysis demonstrated the presence of tight junctions between the epithelial cells and apical microvilli on the surface. Electron dense glycocalyx was detected on the surface of microvilli, indicating the formation of a barrier between the cells on the apical surface and the surroundings (**Figure S2**). Both microvilli and glycocalyx are found on the surface of the conjunctiva and are believed to provide the framework that supports and bind tears, mucus, and immunoglobulins, that have the common function of protecting the eye (Nichols et al., 1983).

Together these findings demonstrate the establishment of the ALI conjunctival epithelium comprised of epithelial progenitors, fibroblasts, basal and superficial conjunctival cells, which express the typical conjunctival specific mucins and show the ultrastructural features of normal conjunctival epithelial tissue in humans. Importantly, these findings validate the suitability of the human conjunctival epithelium ALI model for modelling SARS-CoV-2 infection.

### Conjunctival epithelial ALI organotypic cultures are permissive to SARS-CoV-2 genome expression, but are resistant to productive infection

To assess the permissiveness of conjunctival epithelial and progenitor cells to infection, conjunctival ALI organotypic cultures were inoculated with a clinical isolate of SARS-CoV-2 (BetaCoV/England/2/2020, multiplicity of infection [MOI]=0.5) at the apical surface for 2 hours, inoculum was removed, and infection was assessed regularly for 72 hours post-infection (hpi). Expression of SARS-CoV-2 nucleocapsid (*N*) gene and subgenomic *N* RNA, were detected in cell lysates from 2-72 hpi, suggesting permissiveness to SARS-CoV-2 entry and genome replication. However, there was no significant increase in viral RNA expression over time (**Figure 3A, B**), in contrast to nasal epithelial ALI cultures (Hatton et al., 2021), suggesting a relative resistance to productive replication. Consistent with this, SARS-CoV-2 subgenomic *N* RNA abundance at 72 hpi was at least two orders of magnitude lower than in nasal epithelium ALI cultures infected at a similar MOI (0.1) (**Figure S3B**). Consequently, although SARS-CoV-2 spike protein (S) was detected by western blot at 72 hpi, the expression was substantially lower than infected nasal epithelium ALI cultures (**Figure 3C**). Immunofluorescence analyses at 48 hpi corroborated expression of S protein in conjunctival epithelial cell types, including the mucin secreting cells (**Figure 3E**). These findings suggested that conjunctival cells did not support productive infection. To address this, we measured the release of infectious particles by plaque assays on superficial washes over time. This analysis showed a continuous decline in infectious particle detection from 2 to 72 hpi, indicating that the conjunctival epithelium ALI cultures did not support productive infection (**Figure 3D**). Consistent with these findings, we were unable to identify virion-like structures by TEM analysis at 48 hpi. Together, these data demonstrate that, whilst conjunctival epithelial cells are permissive to SARS-CoV-2 entry and genome replication, they are unable to support productive infection, extending recent findings in an alternative conjunctival model (Singh et al., 2021).

**Figure 3.**
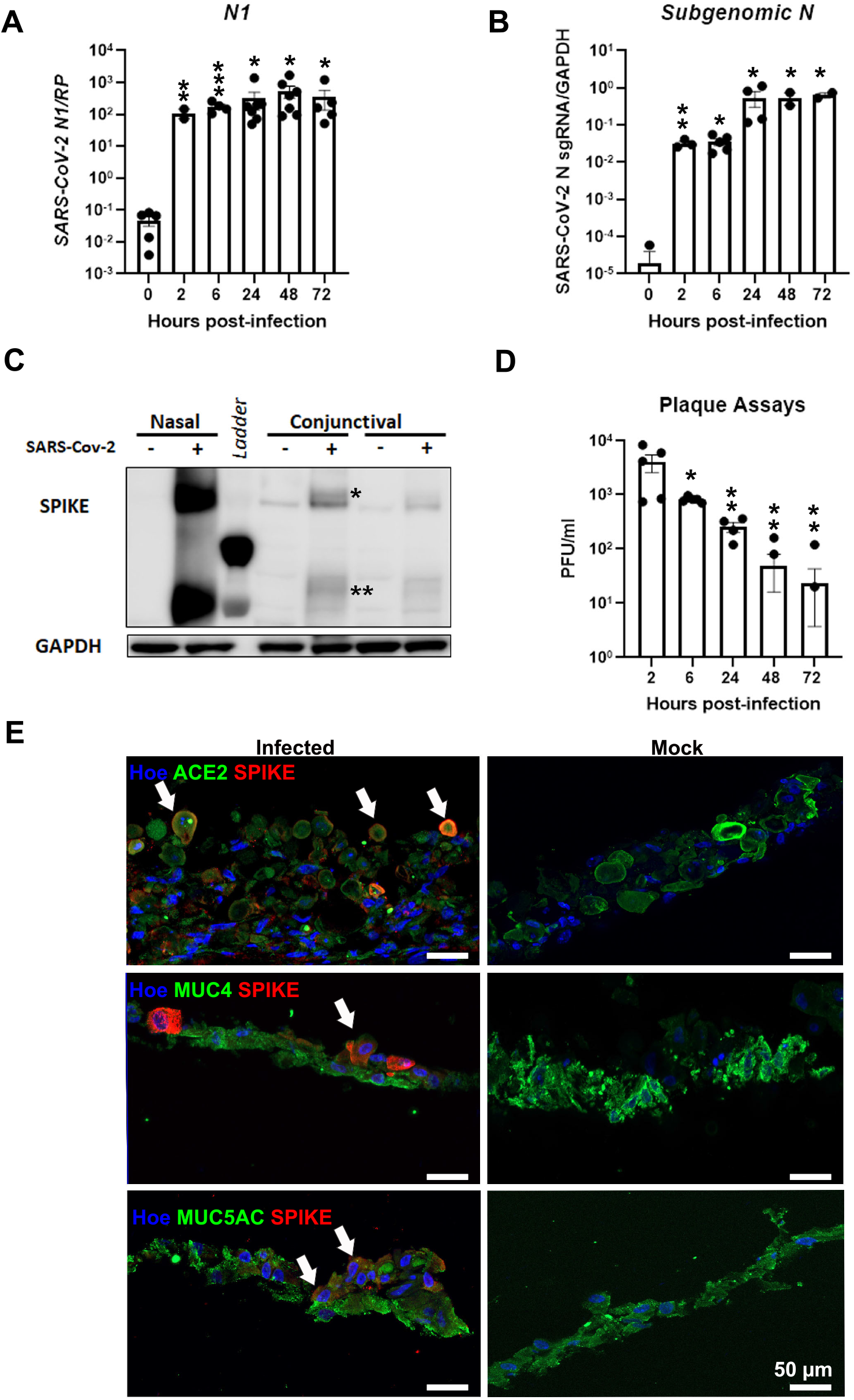
SARS-CoV-2 infection of day 30 human ALI conjunctival organotypic culture. **A, B)** Quantitative RT-PCR expression of nucleocapsid (*N*) gene (normalised to the housekeeper *RNASEP*) and sub-genomic *N* RNA (normalised to *GAPDH*) from 0-72 hpi. Data shown as mean ± SEM, n=3-7 experimental repeats, 3 different donors; * p < 0.05, ** p < 0.01, *** p < 0.001, one way ANOVA with Dunnett’s multiple comparisons to 0 hpi. **C**) Representative western blot showing the expression of SARS-CoV-2 spike (S, shown by *) and cleaved S2 protein expression (shown by **) in the nasal and conjunctival ALI organotypic culture models. GAPDH was used as loading control (representative of repeat experiment in 3 donors). **D**) Release of infectious viral particles was determined by plaque assay using apical washings from 2-72 hpi. Data shown as mean ± SEM, n=2-5, 3 different donors, * p < 0.05, ** p < 0.01, one way ANOVA with Dunnett’s multiple comparisons to 2 hpi. **E**) Immunofluorescence analysis showing the presence of infected cells marked by ACE2 and S co-expression. A few mucin secreting cells are also infected by SARS-CoV-2 as shown by co-expression of MUC4 and MUC5AC with S (white arrows). A panel of mock infected cells is shown on the right-hand side panel, white arrows indicate MUC5AC secreting cells (representative of repeat experiment in 3 donors). Hoe-Hoescht.

### Transcriptional response of conjunctival epithelial cells to SARS-CoV-2 infection

To further assess SARS-CoV-2 cell-type specific tropism, we performed scRNA-Seq at 24 hpi, which represents an early stage in the infection process and a peak of viral gene expression. 15,821 cell transcriptomes from three infected and three uninfected ALI organotypic cultures were integrated and analysed. This analysis defined four major clusters, namely superficial conjunctival epithelium (59.4%), basal conjunctival epithelium (14.3%), epithelial progenitors (25.4%) and fibroblasts (0.9%) (**Figure 4A, Table S2**). To assess relative cell type permissiveness to SARS-CoV-2 infection, viral gene expression was investigated. This analysis showed viral transcripts in all cell types, albeit at low percentage (**Figure 4B, C**). Notably fibroblasts displayed the highest permissiveness with 14.6% of total fibroblasts expressing any viral transcript(s), followed by superficial epithelium (9.6%), basal epithelium (9.2%) and conjunctival epithelial progenitors (8.2%, **Figure 4D**). This was lower than nasal epithelial cells, where viral transcripts were detected in 25-80% of individual cell types (Hatton et al., 2021) by scRNA-seq. Together these data suggest that SARS-CoV-2 is capable of infecting superficial, basal and conjunctival epithelial progenitor cells, but at low efficiency, consistent with our previous findings.

**Figure 4.**
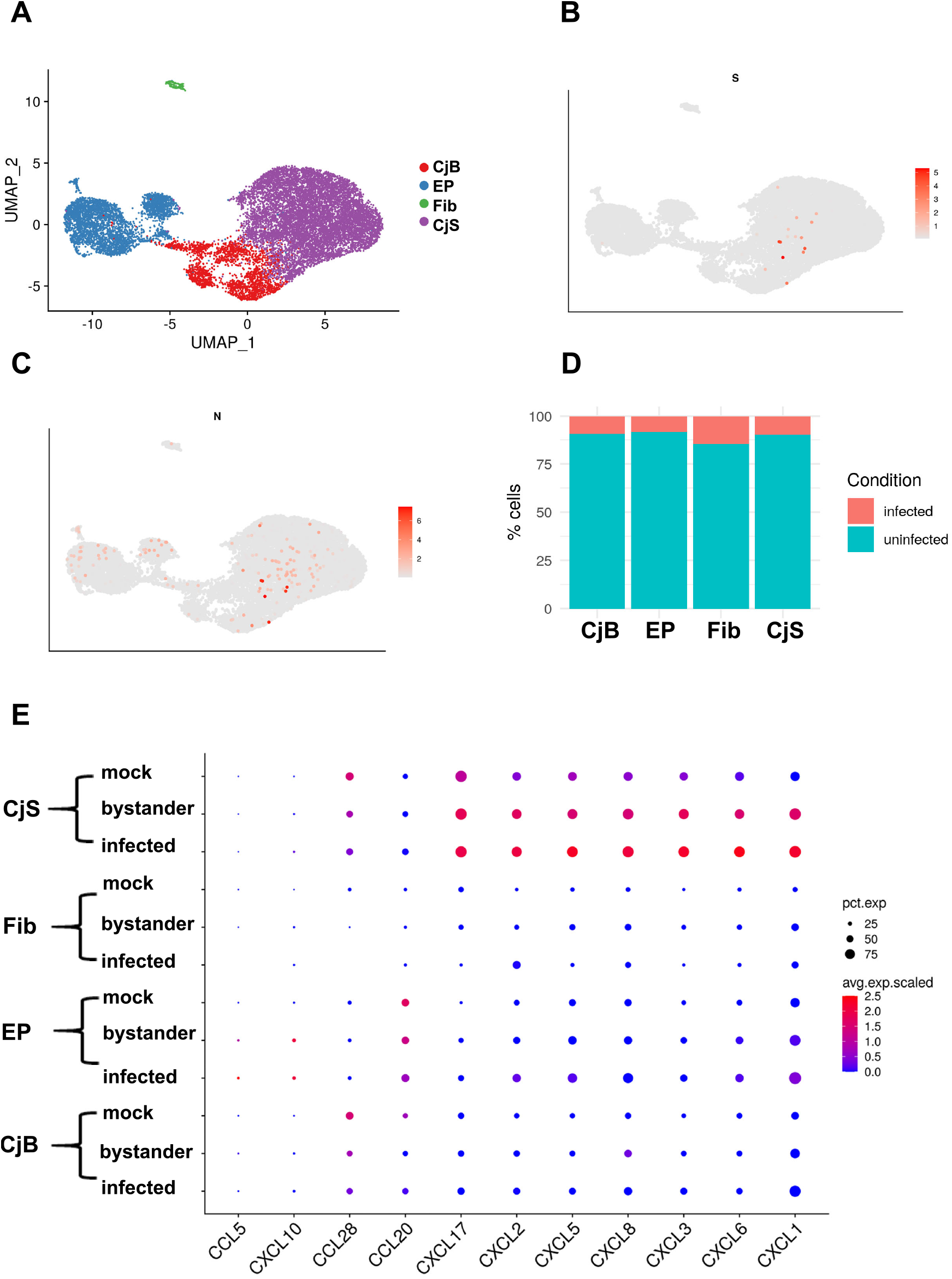
Single cell RNA-Seq analyses at 24 hpi reveal broad but low tropism of SARS-CoV-2 in the ALI conjunctival organotypic culture model. **A)** UMAP visualisation of scRNA-Seq data from mock and infected conjunctival ALI cultures (15,821 cells from three different donors, mock and SARS-CoV-2 infected) showing the presence of epithelial progenitors (EP), superficial (CjS) and basal conjunctival epithelium (CjB) and fibroblasts (Fib). **B,C**) Expression of S and N transcripts shown as superimposed single gene expression plots on the UMAP. **D**) Relative proportion of infected cell types (epithelial progenitors (EP), superficial (CjS) and basal conjunctival epithelium (CjB) and fibroblasts (Fib)) based on expression of any viral transcript. **E**) Dot plot demonstrating expression of key chemokine marker upregulated in response to SARS-CoV-2 infection in all cell types with intensity demonstrated by colour and size of the dot representing the proportion of cells expressing the marker.

Given these findings, we next sought to assess the cell-type specific host cell response to SARS-CoV-2 infection, hypothesising that a more robust innate immune response may account for the reduced permissiveness of conjunctival epithelial cells to SARS-CoV-2. We performed differential gene expression (DE) analysis for each cell type (adjusted p < 0.05), defining three experimental conditions: *SARS-CoV-2 infected* (as defined by detectable expression of at least one viral gene), *SARS-CoV-2 exposed but uninfected* (bystander cells) and *unexposed* (mock-infected cells). Differential gene expression analysis of SARS-CoV-2 infected to mock cells (**Table S3**), revealed the significant upregulation of several chemokines (*CXCL1, CXCL6, CXCL3, CXCL5, CXCL8, CXCL2, CXCL17*) in the superficial conjunctival epithelium (**Figure 4E**) and to a lesser extent in the conjunctival epithelial progenitors (*CXCL5, CXCL17, CXCL10, CXCL3, CXCL6, CXCL2, CXCL1, CXCL8*), corroborating recent findings reported by Eriksen et al in SARS-CoV-2 infected scleral cells (2021). Importantly, several additional TNF and IL1 regulated genes (*C3, CD55, CCL5, CD47*) were upregulated in infected epithelial progenitors, consistent with the predicted activation of various pattern recognition pathways and NF-KB dependent signalling pathways by Ingenuity Pathway Analysis (IPA) (**Table S4, Figure 5A**). Similarly, TNF and IL1 regulated genes were observed in SARS-CoV-2 infected superficial conjunctival epithelial cells (**Table S4, Figure 5B**) alongside the upregulation of *IL6*, indicative of a robust NF-κB response in both cell types. To confirm this, ALI cultures were pre-incubated with the IKKβ inhibitor, BI605906 (10 μM), which blocks NF-kB activation, prior to SARS-CoV-2 infection. This significantly reduced the expression of *CXCL8* and *TNF* (**Figure S4B**), confirming their NF-kB-dependence, yet had no impact on expression of SARS-CoV-2 nucleocapsid (*N*) gene at 24 hpi (**Figure S4A**), suggesting that NF-kB did not substantially impact on viral genome expression under these conditions.

**Figure 5.**
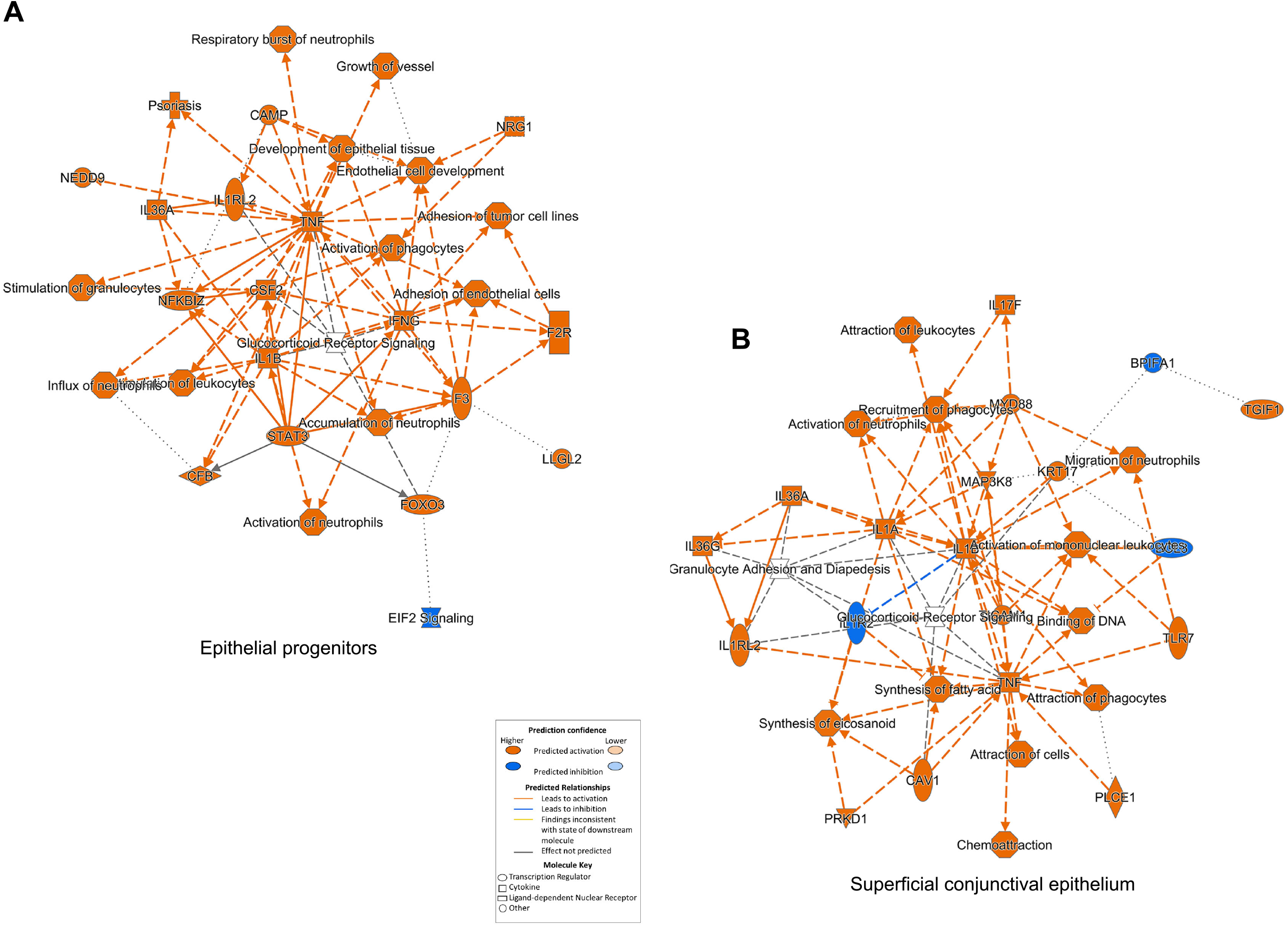
Representative network analysis of predicted regulators in the SARS-CoV-2 infected cells in the epithelial progenitors (A) and superficial cells (B). Differentially expressed genes between infected and mock cells within the epithelial progenitors and superficial conjunctiva epithelium cluster were generated using the Seurat *FindMarkers* function. IPA Upstream Regulator Analysis was used to predict upstream transcriptional regulators from this gene list, using the Ingenuity^®^ Knowledge Base to create mechanistic networks.

Given the prominence of NF-κB driven transcription in infected cells, we undertook gene-set enrichment analysis of NF-κB target genes, finding expression was upregulated in infected cells compared to mock or bystander cells (**Figure 6A**). Notably, there was no widespread induction of antiviral IFN signalling, identified by expression of interferon stimulated genes (ISGs) (**Figure 6B**). Indeed, there was downregulation of certain ISGs, including *IFI6* (also known as IFI-6-16) and *IFI27*-like (also known as ISG12) in superficial conjunctival epithelial cells (**Tables S3, S5**), indicating evasion of an IFN response in infected cells by SARS-CoV-2. Significantly enriched signaling pathways and biological processes in conjunctival superficial and epithelial progenitor cells, included EIF2 stress, glucocorticoid receptor signalling, the coronavirus pathogenesis pathway, complement system, mitochondrial dysfunction and oxidative phosphorylation (**Figure S5**), corroborating recent findings reported by the comprehensive human-SARS-CoV-2 interactome (Kumar et al., 2020).

**Figure 6.**
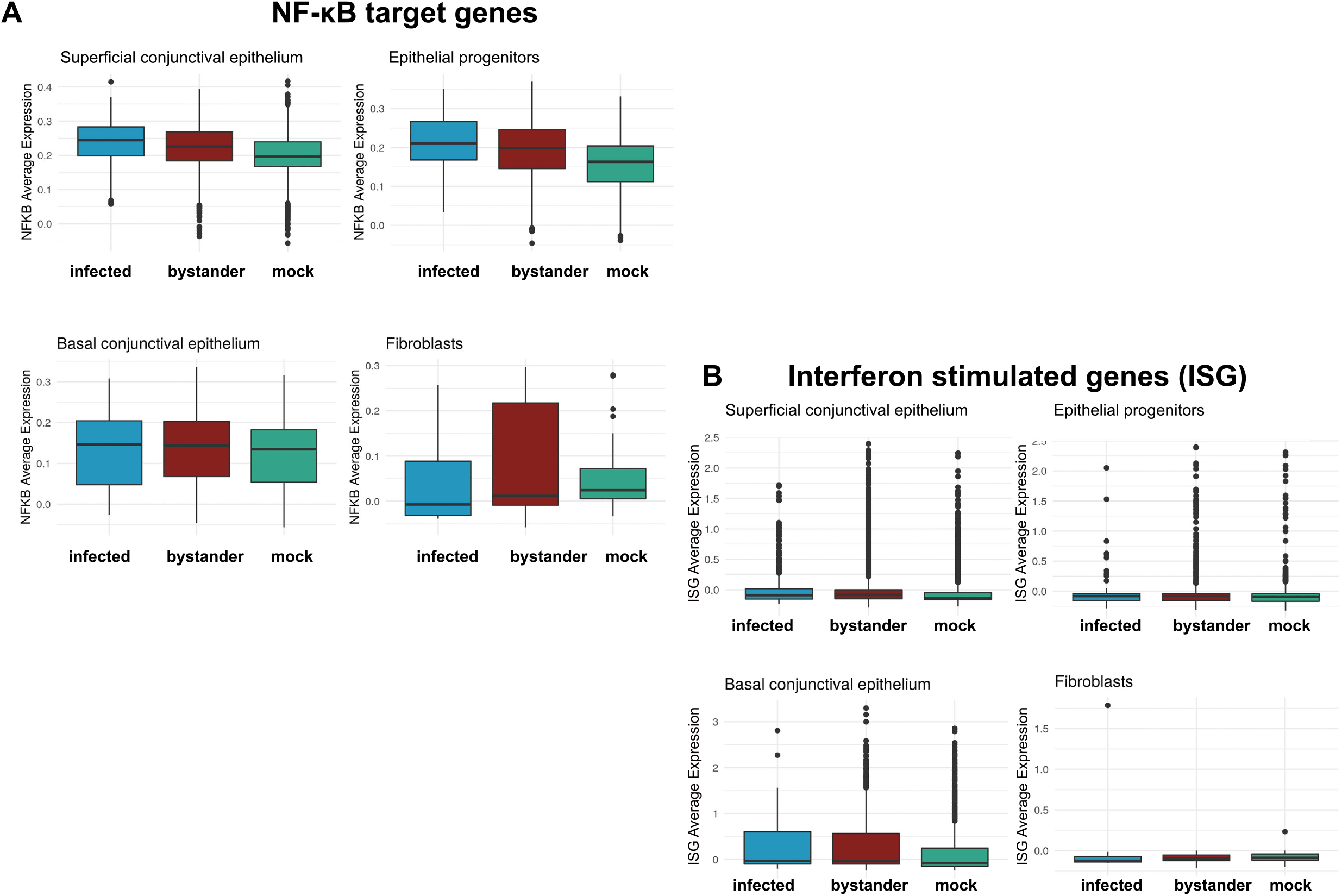
NF-κB target (A) and IFN stimulated gene (ISG) expression (B) in infected, bystander and mock in the epithelial progenitors, superficial and basal conjunctival epithelium and fibroblasts. Gene set scores greater than zero suggest expression levels higher than background gene expression. The bottom and the top of the boxes correspond to the 25th (Q1) and 75th (Q3) percentiles, and the internal band is the 50th percentile (median). The plot whisker minimum is calculated as Q1-1.5 x IQR and the maximum as Q3 +1.5 x IQR. IQR= interquartile range. Outside points correspond to potential outliers.

A more muted transcriptional response to SARS-CoV-2 infection was observed in basal conjunctival cells and fibroblasts with only 19 and 1 differentially expressed genes being identified respectively apart from the viral transcripts (**Table S3**). One chemokine, *CXCL1*, was upregulated when the infected basal conjunctival epithelial cells were compared to mock infected cells, indicating a similar NF-κB response to the epithelial progenitors and superficial conjunctival cells, albeit reduced in terms of repertoire of target gene expression (**Figure 6A**). In addition, *MT1X*, a gene involved in oxidative stress response, was also upregulated (**Table S3**), corroborating reported links between COVID-19 and dysregulation of oxidative stress marker genes (Saheb Sharif-Askari et al., 2021).

A notable feature of SARS-CoV-2 infected epithelial progenitor cells was the upregulation of markers expressed in the superficial conjunctival epithelial cells (*PSCA, PIGR*) and downregulation of basal epithelial markers (*KRT6A, KRT6B*) (**Table S3**), indicating a propensity to differentiate in response to infection, which has not been reported previously and needs to be investigated further. Notably, *SPRR3*, a cornified envelope gene, was significantly upregulated in infected conjunctival epithelial progenitors, whilst *KRT17* was upregulated in the superficial conjunctival epithelial cells (**Table S3**). An increase in expression of genes involved in keratinization has been reported in the tears collected from COVID-19 patients (Mastropasqua et al., 2021). These findings are interesting and suggest a potential molecular pathomechanism underlying the keratoconjunctivitis reported in some of the COVID-19 patients (Loffredo et al., 2021, Al-Namaeh, 2021, Ozturker, 2021).

### Robust paracrine signalling in response to SARS-CoV-2 infection

We next asked whether there was evidence of a paracrine immune signalling response to SARS-CoV-2 in uninfected bystander cells, which are exposed to factors produced by SARS-CoV-2 infected cells but are not themselves infected. The analyses of bystander versus mock-infected cells revealed several DE genes in the conjunctival epithelial progenitors, superficial and basal cells (**Table S5**), suggestive of a robust paracrine response. In general, this response mirrored that of infected cells, in that it was dominated by NF-KB signalling, without evidence of induction of a paracrine antiviral IFN response. Several chemokines were upregulated in bystander conjunctival superficial epithelial and progenitor cells (**Figure 4E**), consistent with predicted activation of upstream regulators including IL1 and TNF (**Figure 7A, B, Table S5**). These data indicate that SARS-CoV-2 infection triggers proinflammatory NF-κB signalling in bystander conjunctival superficial epithelial and progenitor cells, corroborated by activation of NF-κB target genes in bystander versus mock infected cells (**Figure 6A**). Assessment of context specific interferon stimulated gene (ISG) expression identified no evidence of induction of an IFN response in either infected or bystander conjunctival cell types (**Figure 6B**).

**Figure 7.**
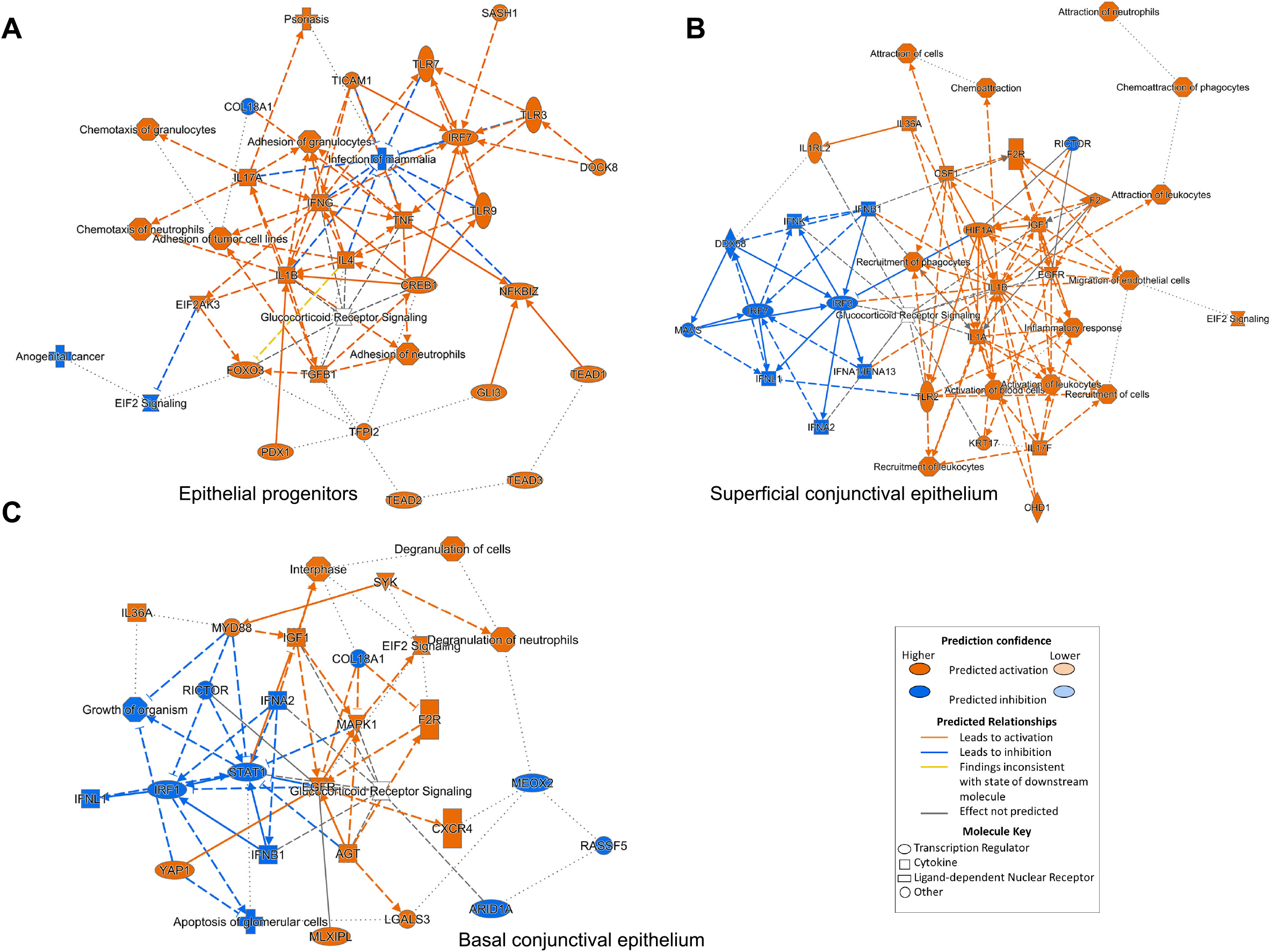
Representative network analysis of predicted regulators in the bystander cells in the epithelial progenitors (A), superficial (B) and basal epithelial cells (C). Differentially expressed genes between bystander and mock infected cells within the conjunctival epithelial progenitors, and superficial and basal conjunctiva epithelium cluster were generated using the Seurat *FindMarkers* function. IPA Upstream Regulator Analysis was used to predict upstream transcriptional regulators from this gene list, using the Ingenuity® Knowledge Base to create mechanistic networks.

Similarly to infected conjunctival epithelial progenitor cells, bystander epithelial progenitors also demonstrated an upregulation of genes expressed predominantly in the superficial conjunctival epithelium (e.g., *PIGR, PSCA, LYPD2, AGR2*) and downregulation of genes expressed in the basal conjunctival epithelium (*KRT6A, KRT6B* etc), indicating a propensity of bystander epithelial progenitor cells to differentiate towards superficial conjunctival epithelium upon exposure to factors produced by infected cells. Expression of genes involved in keratinisation (*SPRR3* in conjunctival epithelial progenitors and *KRT17* in the superficial and basal conjunctival epithelium, **Table S5**) was also observed in bystander cells, indicative of a wider keratinisation occurring in the conjunctival epithelium upon SARS-CoV-2 infection.

Notably, in both superficial and basal conjunctival epithelial cells, there was some evidence of suppression of paracrine type I and III IFN signalling (**Figure 7B, C**), extending recently published findings (Xia et al., 2020, Park and Iwasaki, 2020, Fung et al., 2021, Decker, 2021), suggestive of viral evasion of IFN signalling in bystander cells. It is not clear whether this is due to a paracrine effect of SARS-CoV-2 itself, or an indirect consequence of other host signals, such as TNF, which is recognised to suppress IFN responses in certain contexts (Banchereau et al., 2004). In addition, paracrine activation of epidermal growth factor receptor (EGFR) was predicted in both cell types (**Figure 7B, C**). EGFR is expressed in the plasma membrane of corneal and conjunctival epithelial layers: its expression decreases as cells differentiate. Stimulation with EGF accelerates wound healing, while application of EGFR inhibitors as anti-cancer therapies results in increased risk of corneal erosions (Peterson and Ceresa, 2021). Together, these data suggest an initiation of a “wound like response” in bystander basal and superficial conjunctival epithelial cells.

In both infected and bystander basal and superficial conjunctival epithelial cells, activation of glucocorticoid receptor signalling was also predicted by IPA analyses (**Figures 5, 7**). Glucocorticoids (GCs) are steroids produced and released by the adrenal gland in response to stress (Sulaiman et al., 2018). Topical administration of GCs is used to manage inflammatory insults in the ocular surface, including allergic conjunctivitis and post-surgical inflammation (Friedlaender, 1998). It is thought that GCs supress the expression of pro-inflammatory cytokines, induce the expression of mitochondrial reactive oxygen species and increase apoptosis of human corneal epithelial cells (Ryu et al., 2017). Other findings suggest that GCs receptor activation increases the expression of ocular surface mucins, which may comprise a novel mechanism underlying the therapeutic benefits of GCs in ocular surface inflammatory diseases besides the well-documented anti-inflammatory effects (Taniguchi et al., 2017)_1_ The co-enrichment of pro- (NF-κB) and anti-inflammatory (GCs) pathways suggests a well-balanced response of conjunctival epithelial cells to infections, avoiding hyperproduction of inflammatory cytokines, which merits further investigations.

### Cell type specific transcriptional differences between the bystander and SARS-Cov-2 infected cells

Finally, we compared DE host genes between infected and bystander cells to gain additional insight into the host-virus interaction within infected cells. In superficial epithelial cells, we identified four significantly DE genes, two C-X-C Motif Chemokine Ligand (*CXCL5* and *CXCL6*), Solute Carrier Family 26 Member 4 (*SLC26A4*) and Solute Carrier Family 40 Member 1 (*SCL40A1*) (**Figure S6A-D**). CXCL5 and CXCL6 are neutrophil attracting chemokines, produced by lung epithelial cells in response to SARS-CoV-2 infection (Vabret et al., 2020). The CXCL concentration in body fluids has been shown to correlate with the severity of the disease and thus put forward as a useful biomarker for predicting neutrophil infiltration. Accordingly, *CXCL5* knockout in the mouse decreases lung inflammation without diminishing SARS-CoV-2 viral clearance, suggesting CXCL5 as potential target for controlling/restricting damage in epithelial tissues (Liang et al., 2020). The product of the *SCL40A1* gene, Ferroportin, is an iron exporter protein. Thus an increase in ferroportin expression could deplete intracellular iron and suppress viral replication (Liu et al., 2020). Ferroportin is expressed in enterocytes, macrophages and hepatocytes and our recently published scRNA-Seq data also indicates a high expression in the superficial conjunctival epithelium (Collin et al., 2021a). Iron is required for a range of biological processes necessary for viral replication, including nucleic acid replication and ATP synthesis. Since *SCL40A1* is upregulated only in infected cells and not in bystander cells (**Figure S6C**), intracellular iron depletion may be a novel strategy employed by conjunctival epithelial cells to combat SARS-COV-2 infection. In basal conjunctival epithelial cells, *FGD6* (FYVE, RhoGEF And PH Domain Containing 6) (**Table S7**) was significantly upregulated in infected but not bystander cells (**Figure S6E**). FGD6 is expressed in the basal conjunctival epithelium *in vivo* (Collin et al., 2021a), however there is not much knowledge of the role it may play in this tissue. A recent preprint identifies *FDG6* as one of the top DE genes in infected Calu-3 and Vero E6 cells and in the Ad5-hACE2-sensitized mouse model of SARS-CoV-2 infection (Ghandikota et al., 2021). It will be thus of interest to explore further its involvement/function in SARS-CoV-2 infection in basal conjunctival epithelium. In fibroblasts, *U2AF1L4* (U2 Small Nuclear RNA Auxiliary Factor 1 Like 4), *RGP1* (RAB6A GEF Complex Partner 1) and *TRIM16L* (Tripartite Motif Containing 16 Like) expression was increased when SARS-CoV-2 infected were compared to bystander cells (**Table S7**). U2AF1L4 was recently identified as an interacting protein for nucleocapsid N protein (Chen et al., 2021), whilst RGP1 (**Figure S6F**) was one of the top hits in the screen for SARS-CoV-2 essential host factors and pathways required to mediate infection (Schneider et al., 2021). Their upregulation in SARS-CoV-2 infected fibroblasts deserves further investigation. In epithelial progenitors there were no significant DE genes specific for infected cells.

## Discussion

The human ocular epithelium is continuously exposed to infectious droplets and contaminated fomites. Of the three segments that comprise the ocular surface epithelium, corneal epithelium has been shown to be resistant to SARS-CoV-2 infection (Miner et al., 2020), whilst the adjacent limbal epithelium, which harbours the corneal epithelial stem cells, expresses ACE2 and TMPRSS2 at high levels and appears permissive to viral infection (Eriksen et al., 2021). Yet few studies have addressed the permissiveness of the conjunctival epithelium, the largest exposed component of the human ocular surface. A recent study showed that conjunctival epithelium could be infected with SARS-CoV-2, however the ability of this tissue to sustain productive replication, which is highly relevant to its place as a potential entry route for the virus, and the response of individual conjunctival epithelial cell types to infection, remain unresolved (Singh et al., 2021).

Conjunctival and corneal organ cultures and cultured epithelial monolayers have traditionally been used to examine the capacity of virus infection and replication in the human ocular surface. The *ex vivo* organ cultures are limited in numbers and the monolayer cultures do not capture the involvement of ocular surface mucins or the various cell type interactions, which are necessary for understanding the innate immune response in viral infection dynamics. The ALI organotypic conjunctival model reported in this study overcomes these limitations, as it can be generated in large numbers and comprises all the key conjunctival cell types, including mucin secreting cells, which play an important role in the ocular surface defence against viruses (Mantelli and Argüeso, 2008). Importantly, glycocalyx, a layer of glycolipids and glycoproteins, forming a barrier between the apical surface of the ALI model and the surrounding was observed, mimicking the native barrier of conjunctival epithelium. Consistent with our previous single-cell studies showing expression of relevant entry receptors *ACE2* and *TMPRSS2*, in approximately 6.6% of conjunctival epithelial cells *ex vivo* (Collin et al., 2021b), SARS-CoV-2 infection in this ALI conjunctival epithelium model indicated broad but relatively low tropism of SARS-COV-2 for the various cell types (epithelial progenitors, basal and superficial conjunctival epithelium, fibroblasts). Importantly, our data showed no evidence of productive replication in the conjunctival epithelium, consistent with recent findings in the corneal epithelium (Miner et al., 2020). Reasons for this post-entry restriction in different ocular surface cell types remain to be defined, but contrast with permissiveness of the conjunctival epithelium to other respiratory viruses (Belser et al., 2012). These data are consistent with the apparently paradoxical detection of SARS-CoV-2 nucleic acid or protein in post-mortem tissue samples but the relatively infrequent detection of viral RNA in tears or conjunctival swabs of patients with COVID-19. These data are also consistent with the low incidence of conjunctivitis in patients with SARS-CoV-2 confirmed infection and indicate a low risk of SARS-CoV-2 transmission from a conjunctival transplantation.

Using single cell RNA-Seq analysis, we observed an increase in proinflammatory cytokine expression (e.g. *CXC8, CXCL6, CXCL1* etc.) in SARS-CoV-2 infected conjunctival superficial cells and epithelial progenitors, and to a lesser extent in basal epithelial cells. Proinflammatory cytokine expression is driven by the nuclear factor kappa B (NF-κB) signaling pathway, a family of transcription factors, consisting of RelA, RelB, NF-κB1 NF-κB2 and c-Rel homo/heterodimers with RelA or RelB. These are present in the cytoplasm along with inhibitory proteins that are known as inhibitors of NF-κB (IκBs). Virus infection can activate the multi-subunit IκB kinase (IKK) complex, which leads to phosphorylation of IκBs, resulting in nuclear translocation of NF-κB and activation of a plethora of genes in volved in inflammation, immunity, cell proliferation and apoptosis (Oeckinghaus and Ghosh, 2009). A very recent study has shown that ORF3a, ORF7a and N proteins of SARS-CoV-2 act as NF-κB activators, with ORF7a being the most potent NF-kB inducer and proinflammatory cytokine producer (Su et al., 2021). Multiple pieces of evidence point to activation of NF-κB in ocular surface cells upon infection with adenovirus (Rajaiya et al., 2009), influenza A viruses (Belser et al., 2011), and respiratory syncytial virus (RSV) (Bitko et al., 2004). Together these data indicate that the NF-κB activation we and others (Eriksen et al., 2021) have observed is not specific to SARS-CoV-2, but is a general response to viral infection at the ocular surface (Lan et al., 2012). This NF-κB response was also observed in uninfected bystander cells, suggestive of paracrine signalling. Our hypothesis is that this response is due to secretion of proinflammatory cytokines in the infected cells which activate NF-κB signalling in an autocrine manner, but also cause its activation in the bystander cells in a paracrine manner. While proinflammatory cytokines play a critical role in the defence against viruses and other pathogens in general, their hypersecretion can cause tissue damage (Tak and Firestein, 2001). However, we found no evidence that this response was an important determinant of viral permissiveness, since there was no enhancement of infection in the context of NF-κB blockade. Many studies have shown high level of cytokines in COVID-19 patients, with the phenomenon named as cytokine storm (Lange et al., 2021). Glucocorticoids have an inhibitory action on the NF-κB pathway and are often used to reduce the cytokine feedback on NF-κB activity (Ling and Kumar, 2012). It is worth noting that glucocorticoid receptor signalling pathway was amongst the most enriched signalling pathway in the infected and bystander conjunctival epithelial cells. Together these findings, may indicate a balanced cytokine production in response to SARS-COV-2 infection, which is sufficient to activate the host immune response without causing cytokine storm and excessive tissue damage.

A growing body of evidence indicates that similarly to other viruses, SARS-CoV-2 has evolved mechanisms for evading and/or delaying the antiviral effects of type I and III IFNs response (Blanco-Melo et al., 2020). Our single cell RNA-Seq analysis showed no induction of ISGs in neither infected nor bystander cells, suggesting that IFN signalling does not play a role in conjunctival epithelial defence against SARS-CoV-2 infection, at least in the early stages of infection. Furthermore, the expression of two IFN induced genes, *ISG15* and *RSAD2* was not increased up to 72 hpi (data not shown), indicating that this was not simply due to delayed induction, as has been reported in other epithelial cell types (Hatton et al., 2021).

In addition, this analysis also revealed upregulation of superficial conjunctival epithelium marker expression as well as downregulation of basal conjunctival epithelial markers in the infected and bystander conjunctival epithelial progenitors and basal conjunctival epithelium. This may be in part an attempt to preserve the progenitor and proliferating cell pool, by inducing differentiation of infected cells and those exposed to paracrine signals from the former. Finally, the single cell RNA-Seq analysis also showed upregulation of genes involved in cornification (*SPRR3*), which have also been reported in analysis of tears from patients with COVID-19 (Sopp and Sharda, 2021). These findings may also explain rare reports of keratoconjunctivitis in association with SARS-CoV-2 infection. Together these data suggest that SARS-CoV-2 may be associated with dysregulated ocular keratinization, a serious and potentially debilitating problem that is difficult to manage pharmacologically.

In conclusion the data presented herein show that conjunctival epithelium is permissive to SARS-CoV-2 infection, but without evidence of productive viral replication. This study was performed in organotypic models derived from three different patients, with single cell RNA-Seq data obtained from the peak infection interval. Future work should assess changes in transcriptome of each conjunctival cell type in frequent intervals after infection and in a larger number of donors to get deeper insights into the refractory nature of these cell to viral propagation.

## Materials and Methods

### Human tissue donation

Adult human eyes from three female donors of 52, 78, and 80 years old were donated for research following informed consent. All tissue was provided by NHS Blood and Transplant Tissue and Eye Services or the Newcastle NHS Trust following ethical approval (18/YH/04/20). Human tissue was handled according to the tenets of the Declaration of Helsinki and informed consent was obtained for research use of all human tissue from the next of kin of all deceased donors, or patient themselves, who were undergoing exenteration procedures.

### Epithelial progenitor cell expansion

Human conjunctival epithelial cell expansion was performed using methods described previously for limbal epithelial cell expansion (Collin et al., 2021a). In brief, perilimbal conjunctival tissue was minced into small fragments (~ 1mm^2^) and treated with 0.05% trypsin-EDTA solution (Thermo Fisher Scientific, USA) for 20 minutes at 37°C. The resulting cell suspension was removed from the conjunctival pieces and epithelial medium was added to this suspension to inactivate the trypsin solution. This procedure was repeated three times. The pooled cell suspension was centrifuged for 3 minutes at 1000 rpm in Heraeus Megafuge 16R Centrifuge (Thermo Fisher Scientific, USA), the supernatant removed and the remaining cell pellet re-suspended in epithelial medium containing 3:1 mixture of low-glucose DMEM:F12 supplemented with fetal calf serum 10%, penicillin/streptomycin 1% (all Thermo Fisher Scientific, USA), hydrocortisone 0.4 μg/ml, insulin 5μg/ml, triiodothyronine 1.4 ng/ml, adenine 24 μg/ml, cholera toxin 8.4 ng/ml and EGF 10 ng/ml (all Sigma-Aldrich, UK). Cells were counted and assessed for viability using trypan blue exclusion and a haemocytometer. 30,000 viable conjunctival epithelial cells were added to one 9.6 cm^2^ tissue culture well containing the mitotically inactivated 3T3 fibroblasts and placed in a tissue culture incubator at 37°C with a humidified atmosphere containing 5% CO_2_. The medium was exchanged on the third culture day and every other day thereafter. Several days after, cell colonies with typical epithelial progenitor morphology started to appear and were cultured until they became sub-confluent. Following this 3T3 feeder cells were detached and removed using 0.02% EDTA (Lonza, Switzerland), sub-confluent primary cultures were dissociated with 0.5% trypsin-EDTA (Santa Cruz, USA) to single cell suspension and passaged at a density of 6 × 10^3^ cells/cm^2^. For serial propagation, cells were passaged and cultured as above, always at the stage of subconfluence, until they reached passage 3.

### Generation of ALI conjunctival and nasal organotypic culture model

ALI differentiation of epithelial progenitor cells was performed using a method developed for differentiation of lung epithelial basal cells described by Dvijodrski et al. (2021). 250,000 epithelial progenitor cells were detached from feeders as described above and seeded onto matrigel coated 24 well inserts (ThinCerts™, Greiner Bio-one) and fed apically and basally with BEGM Bronchial Epithelial Cell Growth Medium Bullet Kit (Lonza) supplemented with 10μM Y26732 (Sigma Aldrich) and incubated for 48-72 hours until confluent. Once confluent, the apical medium was removed, and the cells were basally with PneumaCult media (StemCell Technologies) for up to 75 days. The nasal organotypic culture model was performed as described in Hatton et al., 2021.

### Infection of conjunctival ALI cultures with SARS-CoV-2

A clinical isolate from Public Health England of SARS-CoV-2 (BetaCoV/England/2/2020) virus was propagated once in Vero E6 cells. The same viral stock was used for all experiments. As SARS-CoV-2 is a Hazard Group 3 pathogen (Advisory Committee on Dangerous Pathogens, UK), all infection experiments were performed in a dedicated Containment Level 3 (CL3) facility by trained personnel.

Infections of conjunctival ALI cultures were performed at day 30 of differentiation as previously described (Hatton et al., 2021). In brief the virus was diluted in DMEM (ThermoFisher Scientific) to achieve a multiplicity of infection (MOI) of 0.5 and added to the apical side. After 2 hours of incubation, the virus containing medium was removed and the cells were fed basally with PneumaCult media. DMEM was used as inoculum for mock infections. A similar procedure was carried out for the infection of nasal ALI epithelial cultures, described in Hatton et al., 2021. The apical washes were collected in 1x phosphate-buffer solution (1xPBS) at 2, 6, 24, 48 and 72 hpi for plaque assays.

### Plaque Assays

Vero E6 cells were seeded onto a 24-well plate at a density of 200,000 cells per well and incubated at 37°C and 5% CO2 overnight. Apical washes collected from ALI infections were thawed and serially diluted in DMEM with 1% FCS (Gibco), then added to cells and incubated for 2 hours before discarding and adding a 2.4% (w/v) microcrystalline cellulose and 2% FCS (mixed 1:1) overlay (Sigma-Aldrich). The assay was incubated at 37°C and 5% CO2 for 70 hours then fixed in 4% PFA for 1 hour. Plates were rinsed in tap water, stained with 0.25% crystal violet for 10 minutes and plaque were counted to calculate plaque forming units per ml.

### Quantitative RT-PCR

RNA was extracted using TRIzol™ reagent (ThermoFisher Scientific), according to the manufacturer’s instructions, and cDNA was generated using GoScript Reverse Transcription System following the manufacturers protocol (Promega). Viral RNA was detected using the CDC 2019-Novel Coronavirus Real-Time RT-PCR Diagnostic Panel as per Centre for Disease and Control’s optimised protocol (Integrated DNA Technologies) for use with Go-Taq 1-step RT-qPCR Master Mix (Promega). Gene expression profile and SARS-CoV-2 N subgenomic RNA expression were determined with a standard qPCR cycle (50°C for 2 minutes, 95°C for 10 minutes, then 40 cycles of 95°C for 15 seconds, 60°C for 1 minute) using Go-Taq qPCR Master Mix (Promega) on a QuantStudio™ 7 Flex Real Time PCR System (ThermoFisher Scientific). Primer sequences can be found in **Table S8**. Data was interpreted using the ΔCT method.

### Statistical Analysis

Statistical analysis was done using GraphPad Prism (version 9.0.0 121). Data was analysed with Ordinary one-way ANOVA using Tukey’s multiple comparisons test unless otherwise indicated. Graphs are presented as mean ± SEM and are in Log10 format. * p < 0.05 ** p < 0.01 *** p < 0.001 **** p < 0.0001 ns – not significant.

### Transmission Electron Microscopy (TEM)

Infected and mock samples were fixed overnight at 4°C in 2% glutaraldehyde (TAAB Lab Equipment) with 0.1 M sodium cacodylate (pH 7.4). The samples were then secondary fixed in 1% osmium tetroxide (Agar Scientific). Dehydration of samples was achieved with graded acetone (25, 50, 75, 100%) for 30 minutes each. Samples were impregnated with resin at the same graded concentrations up to 75% for 60 minutes each. A final incubation with 100% resin (minimum of 3 changes) for 24 hours at 60°C concluded the embedding process. Sections were taken at 70nm using a MT-XL ultramicrotome and mounted on a pioloform-filmed copper grid. Samples were stained in 2% aqueous uranyl acetate and lead citrate (Leica) and imaged on a Hitachi HT7800 transmission electron microscope. Representative micrographs were captured using an EMSIS Xarosa CMOS Camera with Radius software (version 2.1, EMSIS, Germany).

### Immunofluorescence analysis (IF)

Inserts were fixed in 4% PFA for 1 hour and washed three times in 1xPBS. Two protocols were used to prepare tissue for staining. For apical side staining only, ThinCerts™ were divided into four pieces and mounted on SuperFrost Plus™ slides before staining. For apical and basal staining ThinCerts™ were halved and embedded in OCT (Cellpath) then frozen at −20°C. Blocks were sectioned at 10μm on a Leica CM1860 cryostat to expose both the apical and basal sides of the tissue. Cryosections were dried, washed three times in 1xPBS and blocking solution was applied (10% donkey serum, 0.3% Triton-X in PBS) for 1 hour at room temperature. Primary antibodies (**Table S9**) were applied overnight at 4°C. Slides were washed three times with 1xPBS and secondary antibodies (**Table S9**) were applied for 1 hour at room temperature. AlexaFluora 488 and 546 (ThermoFisher Scientific) were used at 2ug/ml. Hoescht 33342 was added for 10 minutes and slides were washed three times in 1xPBS before mounted with VectaShield (Vector Laboratories). All slides were imaged with an Axioimager Z2 microscope using the Apotome 2 system. Images were taken as Z-stacks and presented as maximum intensity projections (MIPs). Image acquisition was captured through Zen software.

### Western Blotting

Lysis buffer (150mM sodium chloride, 1.0% NP-40 or Triton X-100, 0.5% sodium deoxycholate, 0.1% SDS (sodium dodecyl sulfate), 50mM Tris, pH 8.0)) was applied directly to the apical surface of ALI cultures and cells were removed by gentle scrapping. Protein concentration was determined by BCA Assay (ThermoFisher Scientific). NuPAGE™ reducing agent and LDS sample buffer (ThermoFisher Scientific) were added directly and lysates were heated at 70°C for 10 minutes. 15μg of protein were ran on a Novex™ Bolt 4-12% bis-tris mini gel with NuPAGE™ 1x MES SDS running buffer (ThermoFisher Scientific). 5μl of PageRuler™ Plus Prestained Protein Ladder (ThermoFisher Scientific) was used as a size guide. Proteins were transferred onto PVDF iBLOT 2 Transfer Stacks according to manufacturer’s instructions (ThermoFisher Scientific). Membranes were blocked in 5% non-fat milk with 0.1% tween in PBS (PBS-T). Primary antibody combinations were incubated overnight at 4°C. Membranes were washed in 1xPBS-T and membranes were incubated in secondary antibodies for 1 hour at room temperature. Membranes were washed again in 1xPBS-T and developed with SuperSignal West pico PLUS chemiluminescent substrate (ThermoFisher Scientific) according to manufacturer’s instructions and imaged on an Amersham Imager 600.

### Single Cell (sc) RNA-Seq sample processing

Single cell RNA-Seq was performed on cells harvested 24 hours post infection. ALI cultures were washed apically and basally in 1xPBS and dissociated in 100μl 0.025% Trypsin-EDTA (ThermoFisher Scientific) for 15 minutes at 37°C. Trypsin-EDTA was neutralised, and cells were counted on a haemocytometer to get an optimum concentration of 1000 cells per microliter. Suspensions were centrifuged at 400g for 3 minutes and resuspended in 0.04%BSA/PBS solution. For scRNA-Seq cells were captured and libraries generated using the Chromium Single Cell 3′ Library & Gel Bead Kit, version 3.1 (10x Genomics). scRNA-Seq libraries were sequenced to 50,000 reads per cell on an Illumina NovaSeq 6000.

### scRNA-Seq analysis

The sequencing data was aligned to human reference genome (GRCh38) and the Sars_CoV_2 reference (Ensembl ASM985889v3) using CellRanger Version 3.0.1. The Seurat R library was used for the downstream analysis. The filtered_feature_bc_matrix were imported into R and QC filtering applied. The thresholds were applied to the data were a minimum counts per cell of 2000, minimum genes per cell of 500 and maximum percentage mitochondria of 20%. The DoubletFinder package (McGinnis et al., 2019) was used to find and remove doublets. We performed two integrated analyses of the data. Firstly, the mock samples were combined to study the clustering profiles without infection and secondly the mock and exposed cells were integrated together. Seurat (Butler et al., 2018) was used to normalise the data and the first 2000 highly variable genes, identified through vst selection were chosen for the clustering analysis. The gene expression values were then scaled and the number of counts, number of genes and percentage of mitochondria reads per cell were regressed. A PCA dimension reduction was applied using the selected highly variable genes. This was followed by Harmony (Korsunsky et al., 2019) batch correction where batch was set to sample ID. We then constructed a shared nearest-neighbour graph using the first 10 components of the harmony embeddings. Clusters within the graph using a range of resolutions from 0.2 – 2.2. We chose a resolution of 2.2 which over-clustered the data then applied the FindAllMarkers function and annotated the clusters based on the expression of marker genes as either superficial conjunctival epithelium, conjunctival epithelial progenitors, basal conjunctival epithelium or fibroblasts. The exposed cells were then classified into infected and bystander and differential expression was performed with the following contrasts: infected vs mock; bystander vs mock; and infected vs bystander. ISG and NFKβ target gene scores were generated using the AddModuleScore, which calculates the average expression for each group. The ISG gene list was taken from a published IFN-treated nasal cell dataset (Ziegler et al., 2020), while the NF-KB target gene list was obtained from an online resource (BostonUniversity, 2021). UMAP dimension reduction was performed and the DimPlot and FeaturePlot functions were used to visualise the clusters and expression of genes in individual cells and the DotPlot function was used to visualise the expression of genes within cell types and conditions.

## Acknowledgements

We are grateful to BBSRC UK (#BB/V01126X/1, # BB/T004460/1) for funding this work and Dr. Lucy Clark and Prof. David Steel for the procurement of human eye tissue. We thank Mr Sean Carrie, Dr Jason Powell and Dr Aaron Gardner for provision and characterisation of nasal tissue. Nasal tissue for this study was provided by the Newcastle Biobank which is supported by the Newcastle upon Tyne NHS Foundation Trust and Newcastle University. CJAD is funded by a Wellcome Clinical Research Career Development Fellowship (211153/Z/18/Z) and CFH is supported by a Medical Research Council studentship (MR/NO13840/1). The TEM work was supported by BBSRC (#BB/R013942/1).

## Author contributions

RJ, CH, JSS, MG: experimental design and performance, data acquisition and analysis, contributed to manuscript writing and figure preparation

JC: single cell RNA-Seq data acquisition, fund raising

ES, RH, JMC, MH: single cell RNA-Seq data acquisition and deposition

BV, IH: experimental design and performance, data acquisition and analysis

TD, HM, BW: TEM data acquisition and analysis

PR, MH, SH, CMAK, CW, MB: provided tissue, reagents, expertise, facilities and funding, supervised research

FF, LA: study design, fund raising

RQ, CJAD, ML: study design, data analysis, manuscript writing, fund raising, supervised research

## Conflict of interest

The authors state no conflict of interest.

## The Paper Explained

### Problem

Respiratory viruses including SARS-CoV-2, responsible for the COVID-19 pandemic, enter via the respiratory tract epithelium, however they can also use the eye surface as an additional entry point. The conjunctiva has a large unexposed surface area with mucus producing cells. Clinical reports have shown that some of COVID-19 patients suffer from conjunctivitis in the early stages of infection. To date it is not fully established if the conjunctiva can be used as an additional entry and propagation portal for SARS-CoV-2.

### Results

To assess whether conjunctiva can be infected by SARS-CoV-2, we generated an *in vitro* model by culturing human conjunctival cells under special conditions, which promote cell differentiation and layer stratification through air-liquid contact. In this manuscript, we show that the *ex vivo* model contains all cells found in the native conjunctiva and moreover expresses the two key viral entry factors, ACE2 and TMPRSS2. Infection of the model with SARS-CoV-2 shows that while all conjunctival cells can be infected with the virus, new viral particles that are necessary for propagation cannot be generated. Importantly, the conjunctival cells responded to viral infection by upregulating a key signalling pathway, NF-κB, which is often the first line of defence upon viral infection of eye cells.

### Impact

Our data indicate that the risk of SARS-CoV-2 transmission from a conjunctival organ transplant is low. The lack of viral propagation in conjunctiva should also be informative on provision of protective personal equipment used by medical staff and carers of COVID-19 patients.

## Data availability

The Single cell RNA-Seq data datasets produced in this study are under submission at the Gene Expression Omnibus (GEO).

## Supplementary Information

**Figure S1.**
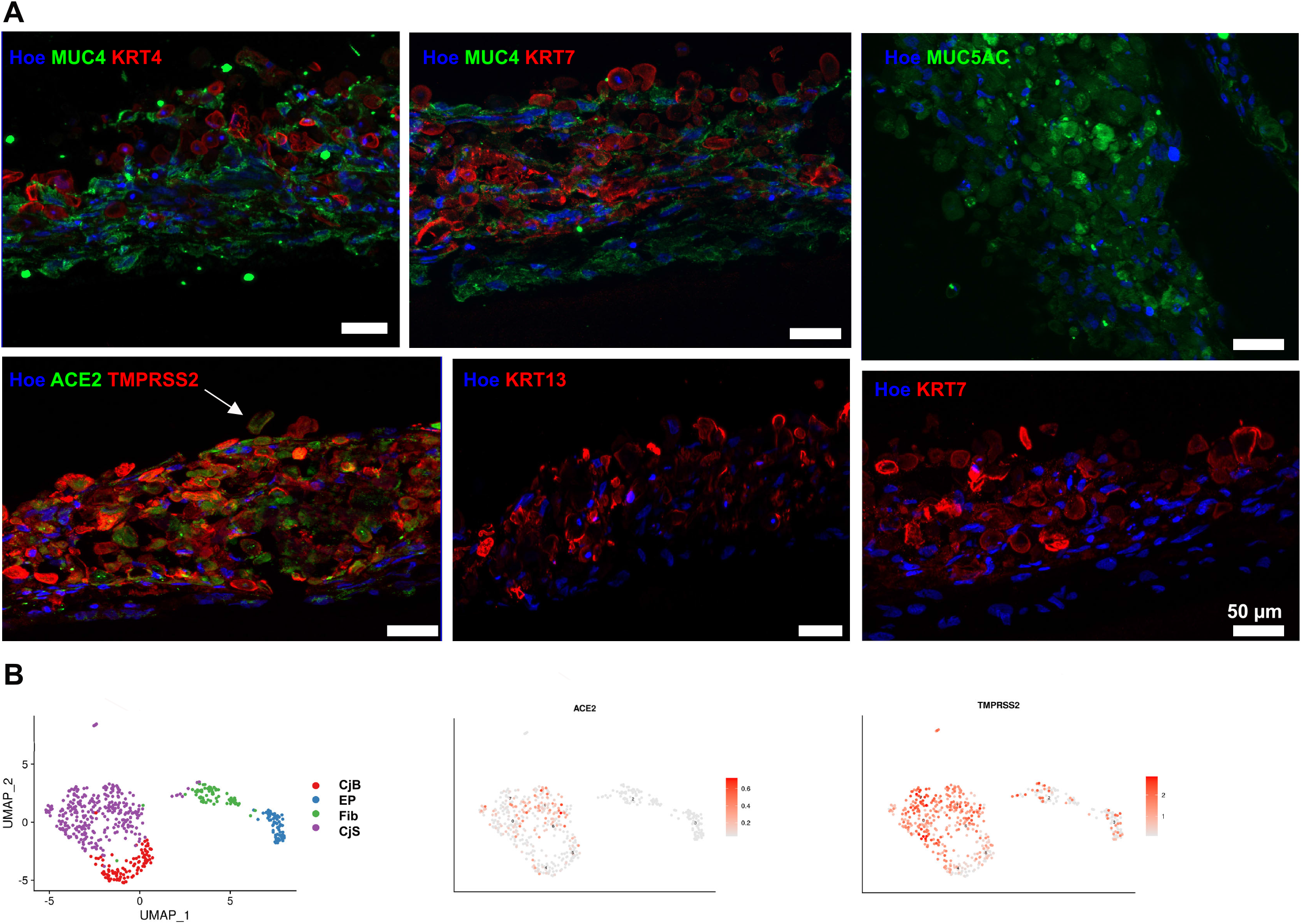
Characterisation of ALI conjunctival organotypic culture model at day 75 of differentiation by immunofluorescence and single cell RNA-Seq. **A)** Immunofluorescence analysis showing coexpression of ACE2 and TMPRSS2 (white arrow) in the superficial layer of the ALI conjunctival organotypic model. KRT7, KRT4 and MUC5AC were predominantly located in the superficial layer, while MUC4 was detected throughout (representative of repeat experiments in three different donor conjunctival ALI cultures). Hoe-Hoescht. **B**) UMAP visualisation of scRNA-Seq data from conjunctival ALI cultures (534 cells from one donor) showing the presence of epithelial progenitors (EP), superficial (CjS) and basal conjunctival epithelium (CjB) and fibroblasts (Fib). Expression of SARS-CoV-2 entry factors, *ACE2* and *TMPRSS2* are shown as superimposed single gene expression plots on the UMAP.

**Figure S2.**
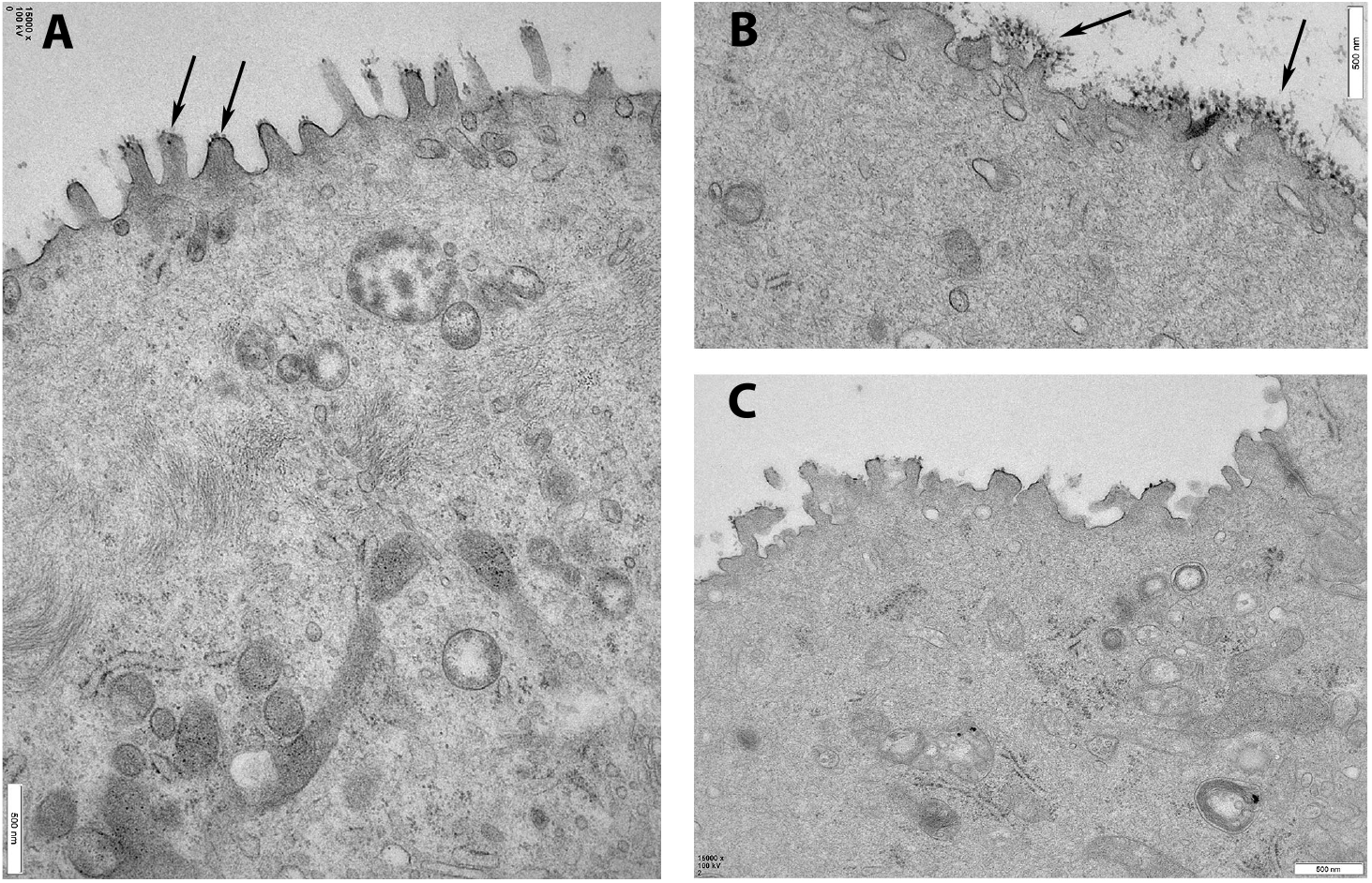
Transmission electron microscopy analysis of ALI conjunctival organotypic culture at day 30. **A**) Numerous apical microvilli are present on the surface of epithelial cells (arrows) indicating cell polarisation. **B**) Fluffy ocular surface-like electron dense glycocalyx (arrows) on surface of microvilli. **C**) Clear tight junctions (arrow) between cells. A-C: representative of repeat experiments in 3 donors.

**Figure S3.**
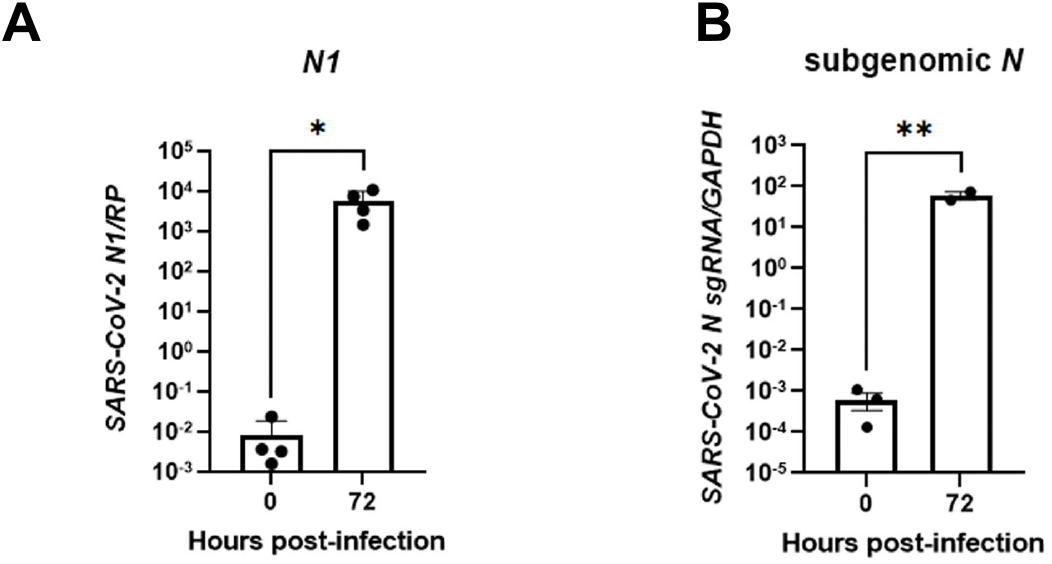
SARS-CoV-2 infection of human nasal ALI organotypic culture. **A, B)** Quantitative RT-PCR expression of nucleocapsid (*N*) gene (normalised to the housekeeper *RNASEP*) and subgenomic *N* RNA (normalised to *GAPDH*) of human nasal ALI organotypic cultures, MOI=0.1. Data shown as mean ± SEM, n=3-4 donors, * p< 0.05 unpaired T-test.

**Figure S4.**
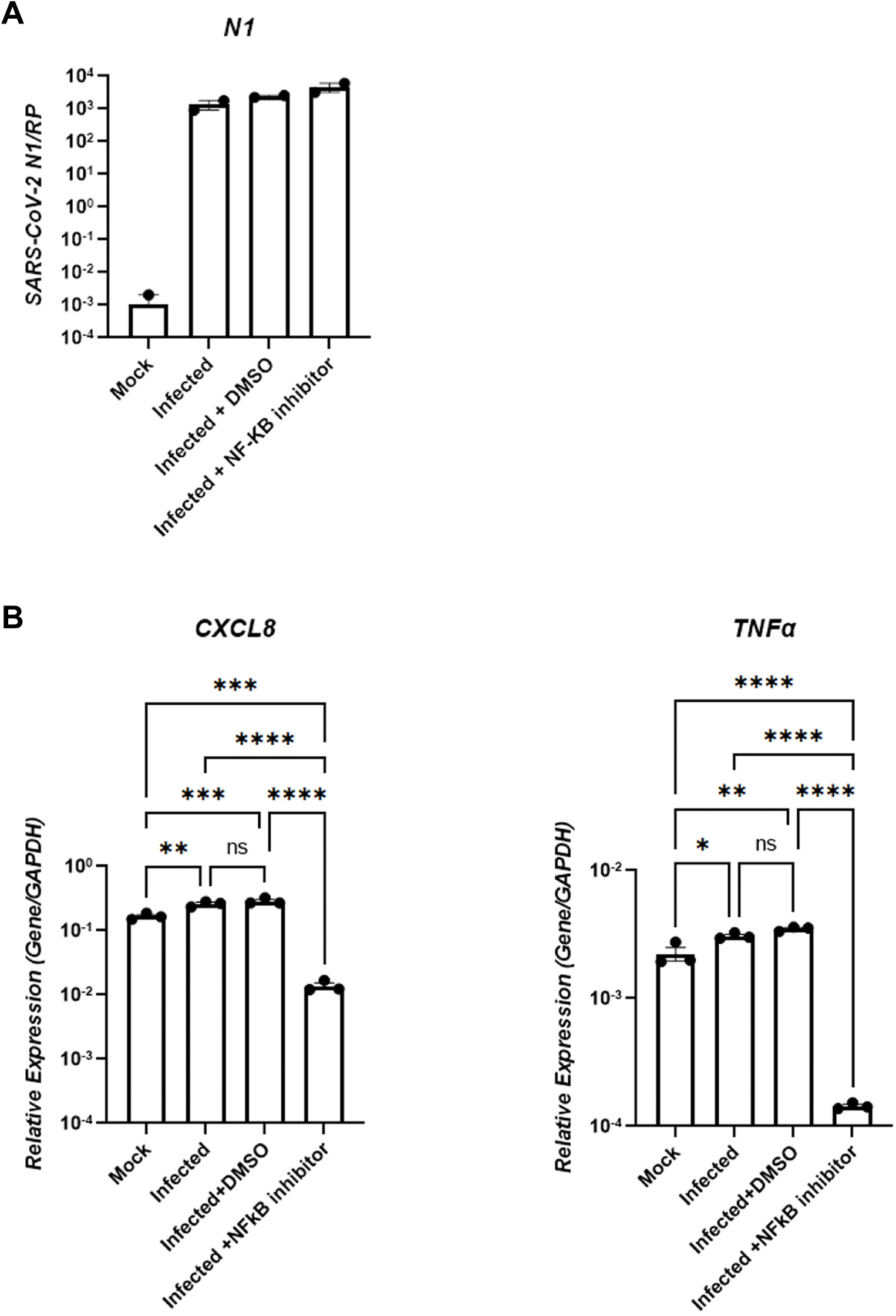
The impact of NF-κB activation on proinflammatory gene expression following SARS-CoV-2 infection of the ALI conjunctival model. **A**) Quantitative RT-PCR expression of nucleocapsid (*N*) gene (normalised to the housekeeper *RNASEP*) at 24 hpi. Data shown as mean ± SEM, n=2 experimental repeats from one donor. The NF-κB inhibitor was diluted in DMSO, hence a DMSO control was included. **B**) Quantitative RT-PCR expression of *CXCL8* and *TNFα* at 24 hpi. Data shown as mean ± SEM, n=3 experimental repeats from one donor. * p < 0.05, ** p < 0.01, *** p < 0.001, **** p < 0.0001, one way ANOVA with Tukey’s multiple comparisons.

**Figure S5.**
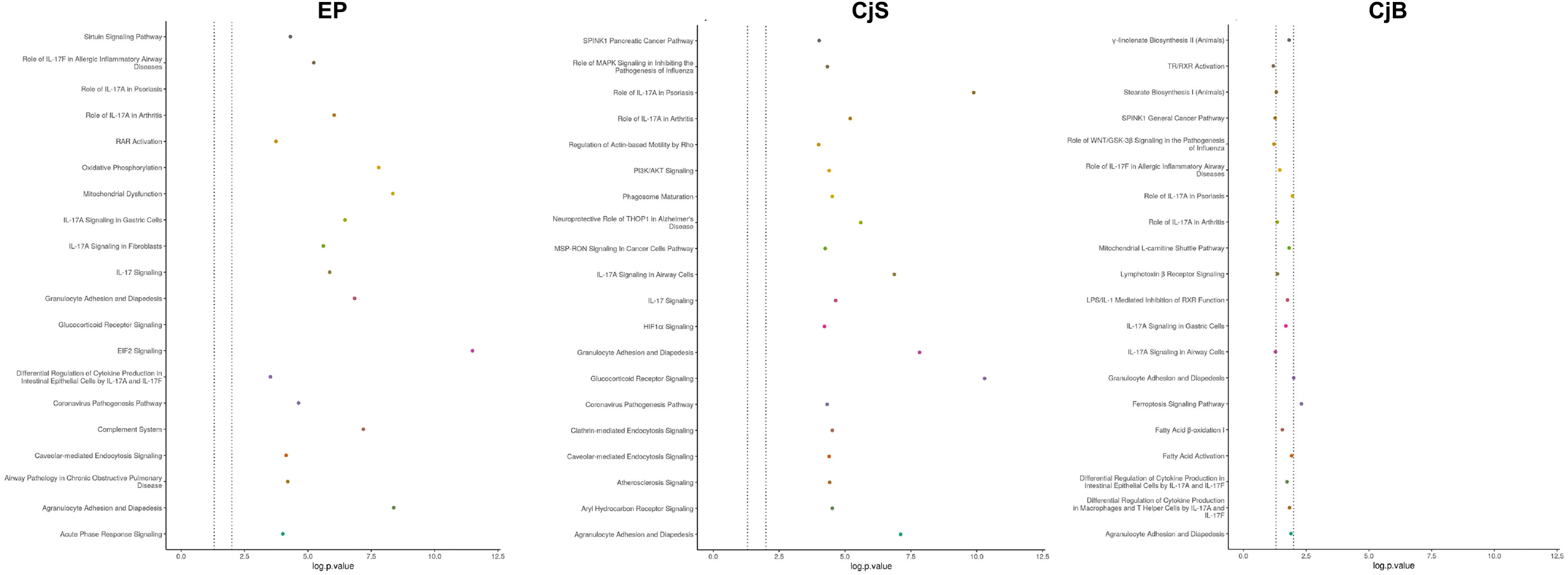
Significantly enriched signaling pathways and biological processes in infected conjunctival superficial and basal and epithelial progenitor cells versus respective mock conditions predicted by the IPA. The dotted lines show a p value of 0.05 and 0.01. Epithelial progenitors (EP), superficial (CjS) and basal conjunctival epithelium (CjB).

**Figure S6.**
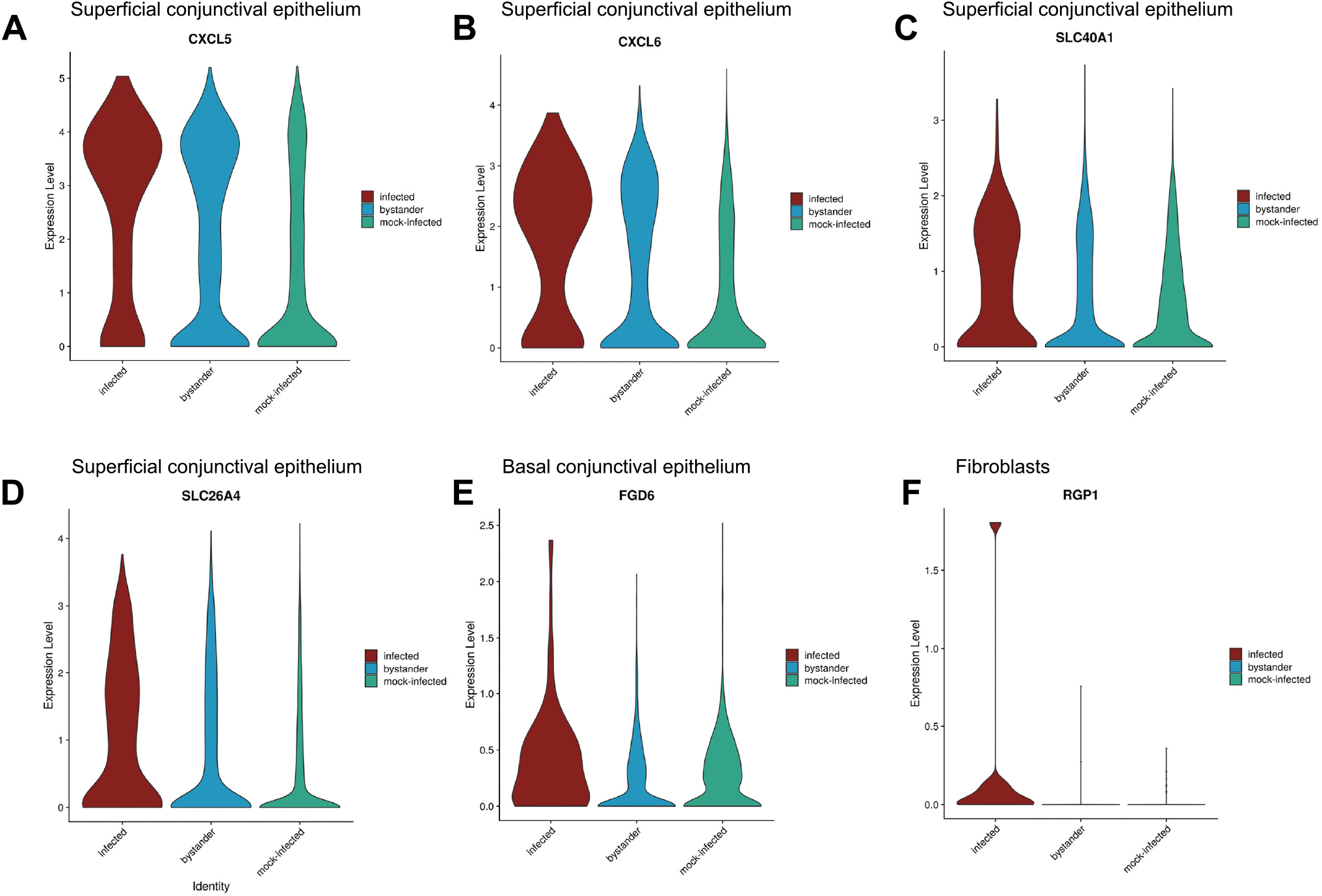
Transcriptional differences between SARS-CoV-2 infected and bystander cells. Expression of genes significantly changes between infected and bystander cells in superficial conjunctival epithelium (**A-D**), basal conjunctival epithelium (**E**) and fibroblasts (**F**) shown as violin plots.

**Table S1**. A full list of highly and differentially expressed genes between the clusters identified in the ALI conjunctival organotypic culture at day 30 and 75 of differentiation.

**Table S2**. A full list of highly and differentially expressed genes between the clusters identified in the SARS-CoV-2 infected and mock ALI conjunctival organotypic culture at day 30 of differentiation.

**Table S3**. A list of differentially expressed genes (p adjust< 0.05) between infected and mock condition in all cell types.

**Table S4**. List of significant regulators of gene expression in the infected cells (versus mock) in all cell types.

**Table S5**. A list of differentially expressed genes (p adjust< 0.05) between bystander and mock condition in all cell types.

**Table S6**. List of significant regulators of gene expression in the bystander cells (versus mock) in all cell types.

**Table S7.** A list of differentially expressed genes (p adjust< 0.05) between infected and bystander in all cell types.

**Table S8**. List of oligonucleotides used in the quantitative RT-PCR analyses.

**Table S9.** A list of antibodies used in this study.

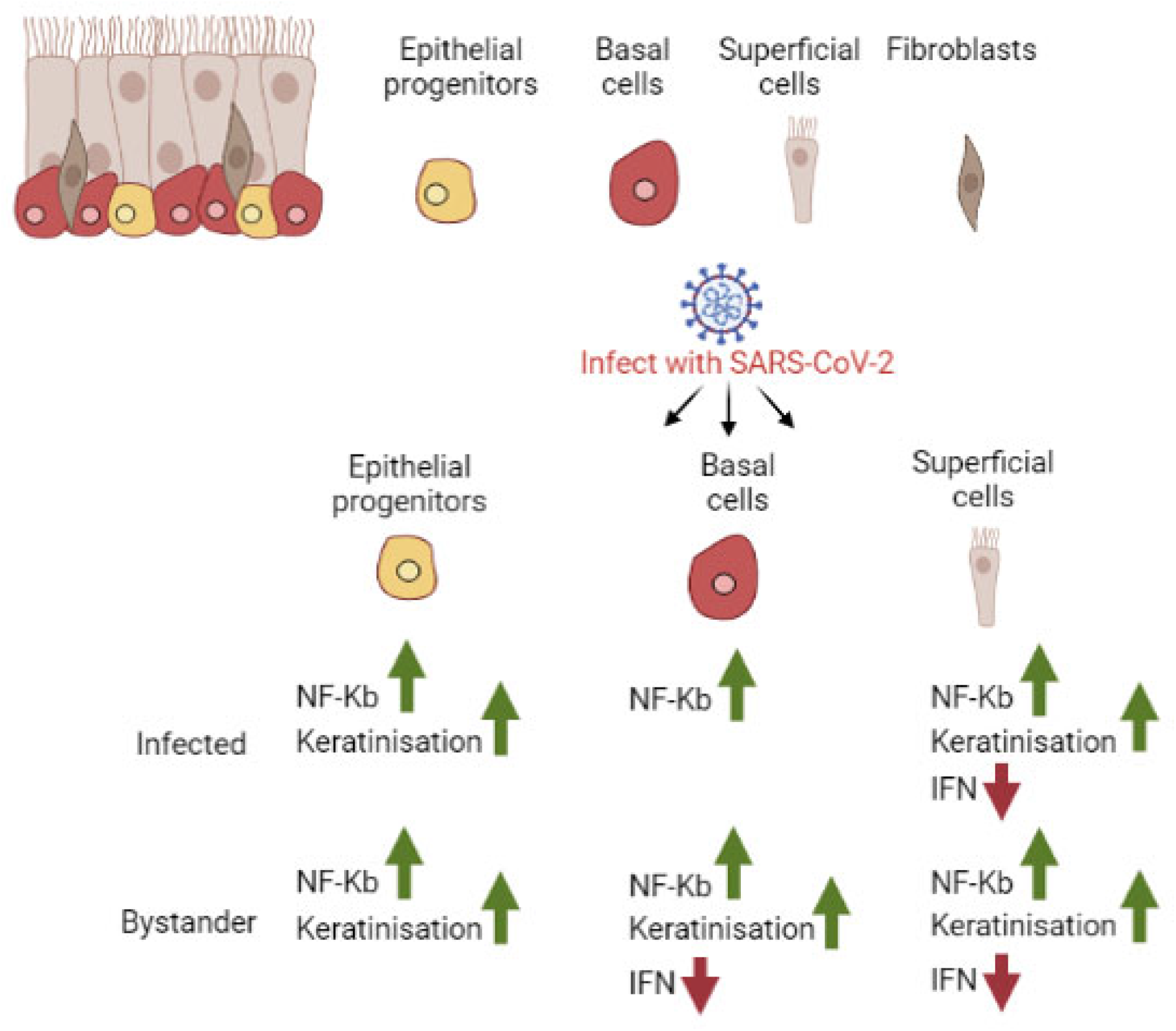

